# *Candida albicans* activates *Staphylococcus aureus* virulence regulatory systems to drive toxin-mediated human cell death

**DOI:** 10.1101/2025.10.03.680095

**Authors:** Kara R. Eichelberger, Nicholas A. Podar, Jia A. Mei, Jacob M. Curry, Ravishankar Chandrasekaran, Saikat Paul, Julie A. Bastarache, Victor J. Torres, Brian M. Peters, James E. Cassat

**Affiliations:** Department of Pediatrics, Division of Pediatric Infectious Diseases, Vanderbilt University Medical Center, Nashville, TN, USA; Department of Pathology, Microbiology, and Immunology, Vanderbilt University Medical Center, Nashville, TN, USA; Department of Host-Microbe Interactions, St. Jude Children’s Research Hospital, Memphis, TN, USA; Department of Clinical Pharmacy and Translational Science, University of Tennessee Health Science Center, Memphis, TN, USA; Division of Allergy, Pulmonary, and Critical Care Medicine, Department of Medicine, Vanderbilt University Medical Center, Nashville, TN, USA; Department of Microbiology, Immunology, and Biochemistry, University of Tennessee Health Science Center, Memphis, TN, USA; Department of Biomedical Engineering, Vanderbilt University, Nashville, TN, USA.; Vanderbilt Center for Bone Biology, Vanderbilt University Medical Center, Nashville, TN, USA; Vanderbilt Institute for Infection, Immunology, and Inflammation (VI4), Vanderbilt University Medical Center, Nashville, TN, USA

**Keywords:** *Candida albicans*, *Staphylococcus aureus*, polymicrobial, SaeRS, human monocytes

## Abstract

Co-infection with *Staphylococcus aureus* and *Candida albicans* leads to worsened disease severity compared to mono-microbial infection. Because our understanding of the mechanisms driving enhanced disease severity during co-infection is limited, we sought to evaluate how interactions with *C. albicans* regulate *S. aureus* virulence towards host cells. We determined that *C. albicans* enhances *S. aureus* cytotoxicity towards murine monocytes via a mechanism requiring the Agr system. Agr is a major regulator of *S. aureus* virulence factors and was previously shown to be activated by *C. albicans,* but the Agr-regulated virulence factors driving immune cell death are unknown. We identified that enhanced murine monocyte cell death requires the α-type phenol soluble modulins and γ-hemolysin. Because several *S. aureus* toxins have species-specific effects, we also tested how co-culture impacts cytotoxicity towards human monocytes. Unexpectedly, we discovered that *C. albicans* induces robust cytotoxicity of an *S. aureus agr* mutant (Δ*agr*), which is completely non-toxic towards murine monocytes. Using reporter strains and combinatorial mutants, we identified that co-culture activates the SaeRS regulatory system in *S. aureus,* and SaeRS is required for human-specific cytotoxicity. We further discovered that the SaeRS-regulated toxin Panton-Valentine Leukocidin (PVL) drives *S. aureus* Δ*agr* cytotoxicity following co-culture. Finally, we observed similar cytotoxicity phenotypes using both *S. aureus* and *C. albicans* clinical isolates, demonstrating broad conservation of this interaction. Interestingly, the magnitude by which *C. albicans* isolates induce cytotoxicity of *S. aureus* Δ*agr* varies among strains tested. Overall, this study identifies that *C. albicans* activates a major *S. aureus* virulence regulatory system in a typically non-toxic strain, triggering *S. aureus* to induce potent human-selective cell death.

## Introduction

*Staphylococcus aureus* is a major human pathogen and a leading cause of multiple invasive infections, including infective endocarditis, osteomyelitis, bacteremia, and pneumonia (1, 2). *S. aureus* infection can be difficult to treat due to the prevalence of antibiotic-resistant strains and high rates of treatment failure, even for infections caused by susceptible isolates (3–6). Colonization is an important risk factor for infection. It is estimated that up to 30% of adults are asymptomatically colonized with *S. aureus* at skin and mucosal sites, such as the gastrointestinal tract, nares, axillae, and groin (7). *S. aureus* is also commonly co-isolated from polymicrobial communities associated with chronic conditions or infections (8, 9). Interactions with other bacteria that are present at colonization and infection sites can play a major role in influencing *S. aureus* virulence and treatment susceptibility (10–15). Fungi are also important constituents of polymicrobial communities that harbor *S. aureus*, but how fungi interact with *S. aureus* to modulate virulence is less well defined. In this study, we sought to determine how interactions with a common co-infecting and co-colonizing fungus, *Candida albicans*, regulate staphylococcal virulence.

The human fungal opportunistic pathogen *C. albicans* is frequently isolated from the same host sites as *S. aureus*. *C. albicans* is among the most often detected species of the mycobiome present in the gastrointestinal tract, the oral cavity, and the vaginal tract of healthy people, and it can also be found on the skin (16–19). The overlap in colonization niches may lead to a greater risk of polymicrobial disease by *C. albicans* and *S. aureus*, especially in patients with co-morbidities or chronic conditions. For instance, *C. albicans* and *S. aureus* are among the two most commonly identified fungal and bacterial species in sputum samples from people with cystic fibrosis (20, 21), burn wounds and diabetic foot ulcers (8, 22, 23), mixed fungal-bacterial bloodstream infection (24), and polymicrobial peritonitis (25). *C. albicans* polymicrobial infections are associated with greater disease severity compared to infection with either bacteria or fungi alone (25–27). A mechanistic understanding of fungal-bacterial interactions that occur among co-colonizing and co-infecting species could inform treatment options to improve disease outcomes.

Murine models of intraabdominal and oral co-infection recapitulate the increased disease severity observed for *C. albicans* and *S. aureus* polymicrobial infections in humans (28–30). In a murine model of polymicrobial intraabdominal infection, *C. albicans* and *S. aureus* co-infection results in higher mortality compared to mono-infection of either organism. Increased mortality is driven by greater activation of the *S. aureus* accessory gene regulator, or Agr, system (31). Agr is a quorum-sensing two-component system that regulates the production of multiple virulence factors in *S. aureus*, including secreted cytolytic peptides and toxins (32, 33). Agr-regulated peptides include the α-type phenol soluble modulins (αPSMs), which exhibit receptor-dependent immunomodulatory effects and receptor-independent cytolytic properties towards host cells (34, 35). Additional secreted toxins regulated by Agr bind specific receptors on host cells, initiating pore formation that induces host cell lysis and contributes to tissue damage during infection (36). Notably, *S. aureus* is a human-adapted pathogen: while some toxins broadly target receptors on cells from multiple host species, several are highly specific for human cells (37–39). Therefore, while murine systems are important for modeling disease, they can only partly recapitulate *S. aureus* toxin-mediated pathogenesis. The importance of Agr-regulated toxins to *S. aureus* pathogenesis and disease severity is underscored by the fact that *S. aureus agr* mutant strains are attenuated in several murine infection models (40–42). Yet, *S. aureus* strains with *agr* mutation or reduced Agr activity are frequently isolated from several human invasive infections or chronic conditions, such as bacteremia, osteomyelitis, and from the lungs of people with cystic fibrosis (43–45). Because many of these chronic infections are polymicrobial, *S. aureus* isolates with Agr dysfunction may still be influenced by interactions with other microbes, including *C. albicans*, at the infection site.

We hypothesized that *C. albicans* enhances *S. aureus* virulence via both Agr-dependent and Agr-independent cytotoxicity towards host cells. To test our hypothesis, we measured cell death of monocytes exposed to mono– and co-cultures. We included both human and murine monocytes to test for species-specific effects of *S. aureus* toxins. Our data demonstrate that *C. albicans* co-culture enhances *S. aureus* Agr-regulated cytotoxicity towards monocytes from both species. Unexpectedly, *C. albicans* also induced cytotoxicity of the typically non-toxic *S. aureus agr* mutant, and this cytotoxicity was selective for human monocytes. We discovered that *C. albicans* activates a second *S. aureus* two-component system, SaeRS, and we determined SaeRS is required for Agr-independent human monocyte death. We further identified that PVL is the main SaeRS-regulated toxin driving induced human cell death. Collectively, this study identifies a novel interaction between *C. albicans* and *S. aureus* that increases virulence selectively towards human cells, demonstrating the importance of investigating *S. aureus* virulence in the context of interactions with polymicrobial communities.

## Results

### *C. albicans* induces *S. aureus* Agr-dependent cytotoxicity towards murine monocytes

Agr-regulated toxins are critical for *S. aureus* initiation of immune cell death. Because *C. albicans* enhances *S. aureus* Agr activation during co-culture (31), we evaluated how *C. albicans* co-culture impacts *S. aureus* cytotoxicity towards monocytes. Monocytes are a major cell type involved in the innate response to *S. aureus* infection and are highly susceptible to multiple Agr-regulated toxins (36, 46). *S. aureus* wild-type (USA300 LAC*) and an isogenic *S. aureus agr* mutant (*agrBDCA::tet,* designated Δ*agr*) were grown either in mono-culture or in co-culture with *C. albicans* wild-type (SC5314). Murine bone marrow-derived monocytes (BMDM) were exposed to culture supernatants, and lactate dehydrogenase (LDH) release was quantified at 6 and 24 hours to determine monocyte cell death. When *S. aureus* wild-type is co-cultured with *C. albicans,* it induces significantly more BMDM cell death (**Fig. 1A-B**). This enhanced cytotoxicity is not simply driven by greater microbial burdens in co-culture, because *S. aureus* and *C. albicans* CFU counts are similar in mono– and co-cultures (**Fig. S1**). *S. aureus* Δ*agr* supernatants were non-toxic towards BMDM, and *C. albicans* co-culture did not induce cytotoxicity of this mutant (**Fig. 1A-B**). These results suggest that *C. albicans* enhances cytotoxicity of *S. aureus* wild-type via Agr-regulated factors, which is congruent with previously published work demonstrating that *C. albicans* enhances *S. aureus* Agr activation during co-culture (31).

**Figure 1.**
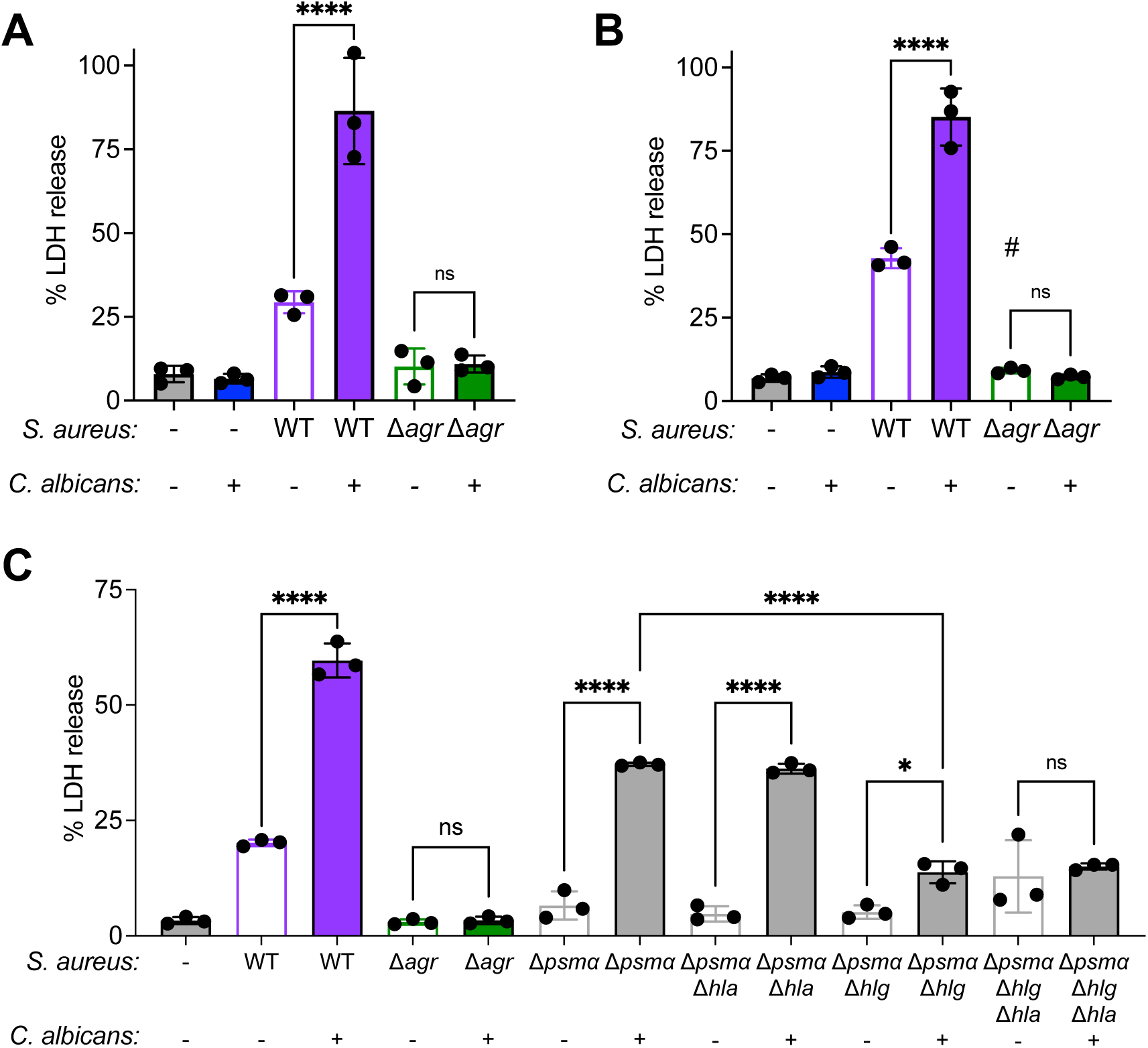
*C. albicans* enhances *S. aureus* cytotoxicity towards murine monocytes via Agr-regulated α-type phenol soluble modulins and γ-hemolysin. Murine bone marrow-derived monocytes were exposed to culture supernatants from *S. aureus* strains grown with or without *C. albicans.* LDH levels in cell culture were quantified at (A) 6 hours and (B-C) 24 hours after supernatant exposure. % LDH release was calculated by standardizing LDH values in each well against cells lysed with 10% Triton-X 100 as a positive cell death control. *S. aureus* WT = wild-type; Δ*agr* = *agrACBD::tet*; Δ*psmα* = *psmα1-4*::*erm*; Δ*hla* = *hla*::*spc*; Δ*hlg* = *hlgACB::tet*. *\**p<0.05, ****p<0.001, and ns (not significant) by one-way ANOVA with Tukey’s multiple comparisons test. # represents p<0.001 compared to *S. aureus* WT mono-culture. N = 3 wells per supernatant condition, error bars = mean±SD.

We next sought to identify the Agr-regulated toxins driving the enhanced cytotoxicity observed during *S. aureus*-*C. albicans* co-culture. We first tested the role of the αPSMs because their expression is directly regulated by Agr, and αPSMs contribute significantly to *S. aureus* cytotoxicity *in vitro* (32, 47). *S. aureus psmα1-4::erm* (designated as Δ*psmα*) mono-cultures were non-toxic to BMDM, indicating that *S. aureus* wild-type mono-culture cytotoxicity is primarily due to the cytolytic activity of αPSMs (**Fig. 1C**). Yet, *C. albicans* induces significant cytotoxicity of *S. aureus* Δ*psmα,* suggesting that other cytolytic factors are involved during co-culture. Since *C. albicans* augments *S. aureus* α-toxin production in an Agr-regulated manner (31), we hypothesized that enhanced α-toxin may be driving enhanced BMDM cell death. However, we did not observe a significant role for this toxin, as *C. albicans* induces cytotoxicity of *S. aureus psmα1-4::erm/hla::spc* (designated as Δ*psmα*/Δ*hla*) towards BMDM (**Fig. 1C**).

Because γ-hemolysin AB (HlgAB) can also induce robust murine monocyte cell death (38), we next hypothesized that HlgAB contributes to co-culture cytotoxicity. *S. aureus psmα1-4::erm*/*hlgACB::tet* (designated as Δ*psmα*/Δ*hlg*) induces minimal BMDM cell death when grown in either mono-or co-culture (5% vs. 13% mean LDH release, respectively), confirming a critical role for HlgAB in driving co-culture cytotoxicity towards BMDM (**Fig. 1C**). The αPSMs are still a major mediator of *S. aureus* cytotoxicity towards BMDM, because loss of γ-hemolysins and α-toxin alone did not impact *S. aureus* cytotoxicity (**Fig. S2**). Collectively, the data demonstrate that *C. albicans* enhances *S. aureus* cytotoxicity towards murine BMDM primarily through Agr-regulated production of αPSMs and γ-hemolysin.

### *C. albicans* induces cytotoxicity of *S. aureus* Δ*agr* towards human monocytes

Because several *S. aureus* toxins selectively target human cells but not murine cells (36), we also tested if *C. albicans* induces *S. aureus* cytotoxicity towards primary human CD14^+^ monocytes. Co-culture induces significantly greater human monocyte cell death at both 6 and 24 hours after supernatant exposure (**Fig. 2A-B**). Cytotoxicity of *S. aureus* wild-type mono-culture is driven by Agr-regulated toxins, because *S. aureus* Δ*agr* is non-toxic towards human monocytes (**Fig. 2A-B**). Surprisingly, *C. albicans* co-culture induces significant cytotoxicity of *S. aureus* Δ*agr* towards primary human monocytes as early as 6 hours after supernatant exposure (**Fig. 2A**). By 24 hours, *S. aureus* Δ*agr* co-culture induces equivalent amounts of human monocyte death as *S. aureus* wild-type mono-culture (**Fig. 2B**). We also confirmed that co-culture induces *S. aureus* Δ*agr-*mediated cell death in human monocytes using kinetic imaging with SYTOX green (**Fig. S3**). Our results demonstrate that in addition to enhancing Agr-dependent cytotoxicity, *C. albicans* is also able to induce *S. aureus* cytotoxicity independently of the Agr system, with specificity for human cells.

**Figure 2.**
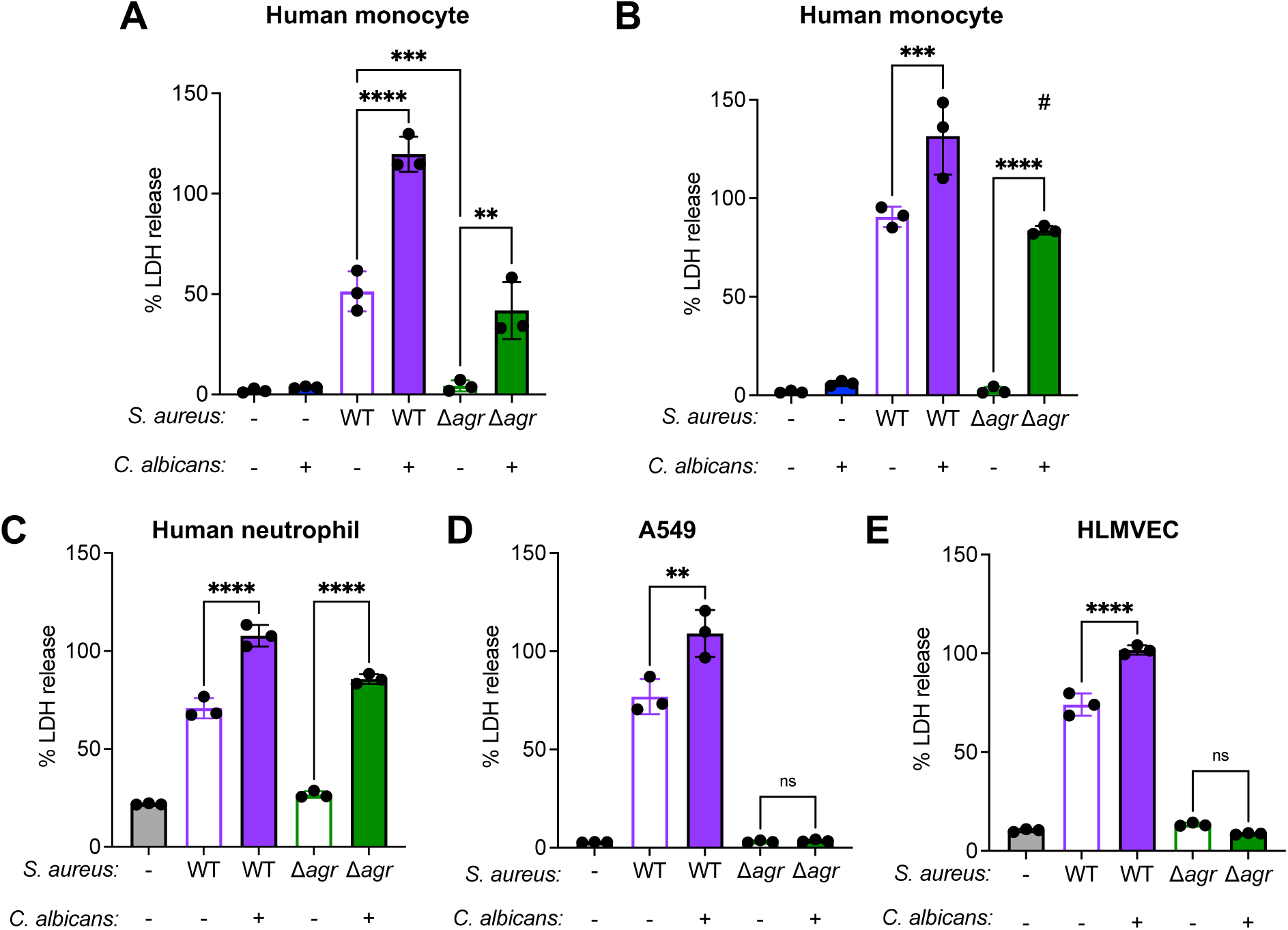
*C. albicans* induces *S. aureus* Δ*agr* cytotoxicity towards human monocytes and neutrophils. (A-B) Primary human CD14^+^ monocytes, (C) primary human neutrophils, (D) A549 human lung epithelial cells, (E) and human lung microvascular endothelial cells (HLMVEC) were exposed to culture supernatants from *S. aureus* cultured with or without *C. albicans.* LDH levels in cell culture were quantified at (A) 6 hours and (B-E) 24 hours after supernatant exposure. % LDH release was calculated by standardizing LDH values in each well against cells lysed with 10% Triton-X 100 as a positive cell death control. *S. aureus* WT = wild-type; Δ*agr* = *agrACBD::tet*. **p<0.01, ***p<0.005, ****p<0.001, and ns (not significant) by one-way ANOVA with Tukey’s multiple comparisons test; n = 3 wells per supernatant condition, error bars = mean±SD.

We next tested if *C. albicans* induces cytotoxicity of *S. aureus* Δ*agr* towards other human cell types. We applied supernatants from *S. aureus* wild-type and Δ*agr* strains cultured with and without *C. albicans* to primary human neutrophils, and we quantified LDH release 24 hours after supernatant exposure. *S. aureus* wild-type mono-culture triggered significant neutrophil cell death, and *C. albicans* co-culture enhanced this cytotoxicity (**Fig. 2C**). *C. albicans* induces significant *S. aureus* Δ*agr* cytotoxicity towards human neutrophils (80% LDH release), like what we observed towards human monocytes (**Fig. 2C**). To determine if *S. aureus* Δ*agr* is cytotoxic to non-immune cells following co-culture, we also tested human epithelial and endothelial cells. *C. albicans* significantly enhances *S. aureus* wild-type cytotoxicity towards A549 human lung epithelial cells (**Fig. 2D**). However, *C. albicans* failed to induce cytotoxicity of *S. aureus* Δ*agr* towards these cells, as both mono-culture and co-culture remained non-toxic (**Fig. 2D**). These results indicate that A549 cells are primarily susceptible to Agr-regulated toxins. We obtained similar results for human lung microvascular endothelial cells (HLMVEC), in that *C. albicans* enhanced *S. aureus* wild-type cytotoxicity but did not induce *S. aureus* Δ*agr* cytotoxicity towards these cells (**Fig. 2E**). Taken together, these data demonstrate that *C. albicans* enhances both *S. aureus* Agr-dependent and Agr-independent cytotoxicity towards human cells. Additionally, human monocytes and neutrophils, but not epithelial and endothelial cells, are susceptible to Agr-independent cytotoxicity.

### *C. albicans* induces Agr-independent cytotoxicity towards human monocytes via *S. aureus* SaeRS

Our discovery that *C. albicans* induces cytotoxicity of a *S. aureus agr* mutant strain towards human monocytes and neutrophils suggests that *C. albicans* induces Agr-independent production of human-specific toxins that target immune cells (36). While Agr is a major regulator of toxin production*, S. aureus* virulence is controlled by a complex and often overlapping network of global transcriptional regulators (48). One such regulator is the SaeRS two-component system, which controls the production of multiple virulence factors including several toxins that demonstrate greater specificity for human cells compared to murine cells (49). SaeRS activity can be indirectly regulated by the Agr system, but several studies demonstrate Agr-independent activity of SaeRS that induces toxin gene expression during *S. aureus* infection (45, 50). Therefore, we hypothesized that *C. albicans* induces SaeRS activation in *S. aureus* Δ*agr,* which in turn mediates greater cytotoxicity towards human cells. To test this hypothesis, we first evaluated cytotoxicity of supernatants from *S. aureus* Δ*agr, saeQRS::spc* (designated Δ*sae*), and an *agrBDCA::tet saeQRS::spc* double mutant (designated Δ*agr*Δ*sae*) cultured with and without *C. albicans*. *S. aureus* Δ*sae* grown in mono– and co-culture induces high levels of human cell death, supporting the important role Agr-dependent toxins in mediating human cell cytotoxicity (**Fig. 3A**). However, while *C. albicans* induces significant cytotoxicity of *S. aureus* Δ*agr,* co-culture fails to induce *S. aureus* Δ*agr*Δ*sae* cytotoxicity towards human monocytes (**Fig. 3A**). These data support that SaeRS is required for *S. aureus* Δ*agr* co-culture cytotoxicity, indicating a critical role for Sae-regulated toxins in inducing human cell death.

**Figure 3.**
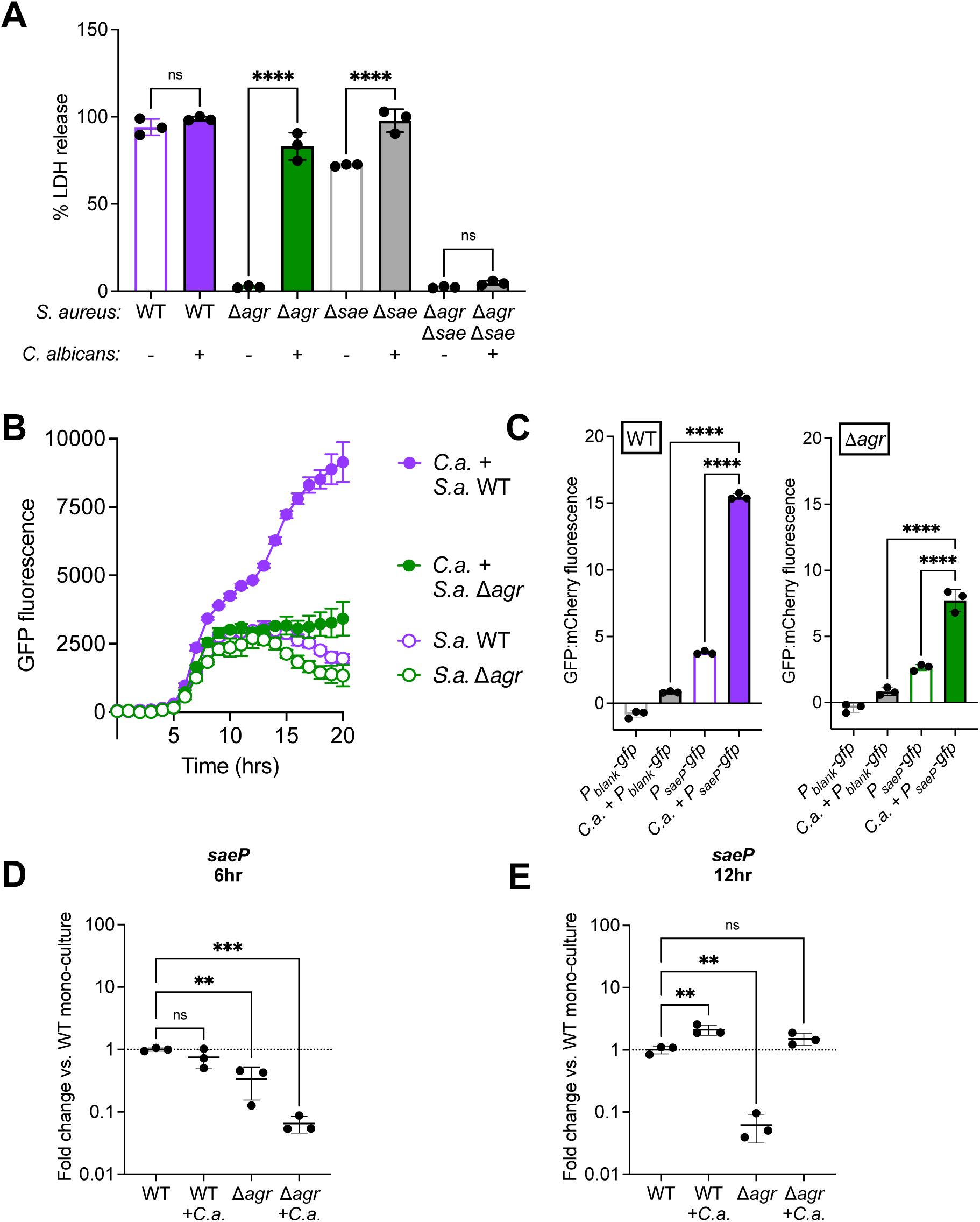
*C. albicans* co-culture induces *S. aureus* SaeRS activation, and SaeRS is required for induced *S. aureus* Δ*agr* cytotoxicity towards human monocytes. (A) Primary human CD14^+^ monocytes were exposed to supernatants from *S. aureus* strains cultured with and without *C. albicans*. % LDH release was quantified at 24 hours after supernatant exposure and calculated by standardizing LDH values in each well against cells lysed with 10% Triton-X 100 as a positive cell death control. (B-C) *S. aureus* wild-type and Δ*agr P_saeP_*-*gfp* or *P_blank_-gfp* strains were cultured with and without *C. albicans,* and GFP and mCherry fluorescence values were measured every hour for 20 hours. (B) GFP fluorescence for each *P_saeP_*-*gfp* strain was blanked against the isogenic *P_blank_-gfp* control fluorescence values. (C) Ratio of GFP was standardized against mCherry fluorescence values for each strain in (B) at 15 hours. (D-E) qRT-PCR analysis of *saeP* transcript levels for cultures at 6 hours (D) and 12 hours (E) of culture. Values are plotted as fold change relative to the WT mono-culture samples. *C. albicans* = *C.a. S. aureus* WT = wild-type; Δ*agr* = *agrACBD::tet*; Δ*sae* = *saeQRS::spc*. For bar graphs, ****p<0.001 by one-way ANOVA with Tukey’s multiple comparisons test; for qRT-PCR, **p<0.01, ***p<0.005, and ns (not significant) by one-way ANOVA with Dunnett’s multiple comparisons test. N = 3 wells or replicates per condition, bars or lines = mean±SD.

To evaluate if co-culture activates *S. aureus* SaeRS, we utilized *S. aureus* wild-type and Δ*agr* strains harboring either a chromosomally-integrated SaeRS-inducible promoter driving *gfp* expression (*P_saeP_-gfp*) or *gfp* without a promoter (*P_blank_-gfp*) as a control (51). All strains also express *mCherry* from a constitutive promoter (P*_sarA_*) (51). *C. albicans* enhances SaeRS reporter activation relative to mono-culture for both *S. aureus* wild-type and Δ*agr* (**Fig. 3B**). Compared to the mono-culture for each strain, activation of SaeRS was about 5-fold higher in wild-type co-culture and 2.5-fold higher in Δ*agr* co-culture. The mCherry fluorescence values were similar among all strains and culture conditions tested (**Fig. S4**). Co-culture significantly enhances SaeRS activation compared to mono-culture for both *S. aureus* wild-type and Δ*agr* at 15 hours, or the time at which culture supernatants are collected to test cytotoxicity (**Fig. 3C**).

To further support that *C. albicans* activates SaeRS in both *S. aureus* wild-type and Δ*agr*, we used qRT-PCR to quantify *saeP* transcript levels in cultures. Expression of *saeP* is significantly reduced in both *S. aureus* Δ*agr* mono– and co-cultures relative to wild-type mono-culture at 6 hours (**Fig. 3D**). By 12 hours, there is about 2-fold greater *saeP* transcript levels in *S. aureus* wild-type co-culture (**Fig. 3E**). Co-culture also increases *saeP* transcript levels in *S. aureus* Δ*agr*, while *S. aureus* Δ*agr* mono-culture expression is significantly lower (**Fig. 3E**). We next tested cytotoxicity of *S. aureus* Δ*agr* mono– and co-culture after 6 and 12 hours of growth, and we observed that co-culture induces significant human cell death after 12 hours of co-culture, but not after 6 hours (**Fig. S5**). The cytotoxicity of these supernatants correlates with when *saeP* transcript levels are increased in *S. aureus* Δ*agr* co-culture. Collectively, these data reveal that *C. albicans* enhancement of *S. aureus* cytotoxicity towards human monocytes requires both Agr– and SaeRS-regulated toxins. Furthermore, *C. albicans* co-culture activates the SaeRS system in *S. aureus* via a mechanism independent of the Agr system.

### *C. albicans* induction of *S. aureus* Δ*agr* cytotoxicity is mediated through Panton-Valentine Leukocidin (PVL)

Our data reveal that co-culture with *C. albicans* induces *S. aureus* Δ*agr* cytotoxicity towards human monocytes but not murine monocytes, and this enhanced cytotoxicity requires *S. aureus* SaeRS. Therefore, we hypothesized that enhanced co-culture cytotoxicity was driven by SaeRS-regulated toxin(s) that exhibit selective activity towards human cells. Previous work from our group identified several *S. aureus* pore-forming toxins that had reduced abundance in the exoproteome of *S. aureus* Δ*sae* compared to *S. aureus* WT, including α-hemolysin, the ɣ-hemolysins (HlgAB and HlgCB), LukAB, LukED, and LukSF-PV (PVL) (47). Of these SaeRS-regulated toxins, LukAB, PVL, and HlgCB induce human cell lysis but have minimal activity towards murine cells (37–39). To test if *S. aureus* Δ*agr* co-culture requires one of these three toxins for human cell death, we examined the cytotoxicity of *S. aureus* double mutants for each toxin in an *agrA* mutant background. Like *S. aureus* Δ*agr*, *agrA::tn* (designated Δ*agrA*) is non-toxic towards human monocytes when grown in mono-culture, and *C. albicans* co-culture induces significant cytotoxicity (**Fig. 4A**). Co-culture induces significant cytotoxicity of Δ*agrA/*Δ*lukAB* and Δ*agrA/hlgACB::tet* towards human monocytes, indicating that neither LukAB nor HlgCB are required for co-culture cytotoxicity towards these cells. *C. albicans* fails to induce cytotoxicity of Δ*agrA/lukSF-PV::spc* towards human monocytes (**Fig. 4A**), demonstrating that PVL is the main toxin required for *S. aureus* Δ*agr* co-culture cytotoxicity. Mono-cultures of *S. aureus* Δ*lukAB, S. aureus lukSF-PV::spc,* and *S. aureus hlgACB::tet* all induce significant human monocyte death (**Fig. S6**). This supports that multiple toxins, including both Agr-regulated and Sae-regulated, contribute to *S. aureus* wild-type cytotoxicity.

**Figure 4.**
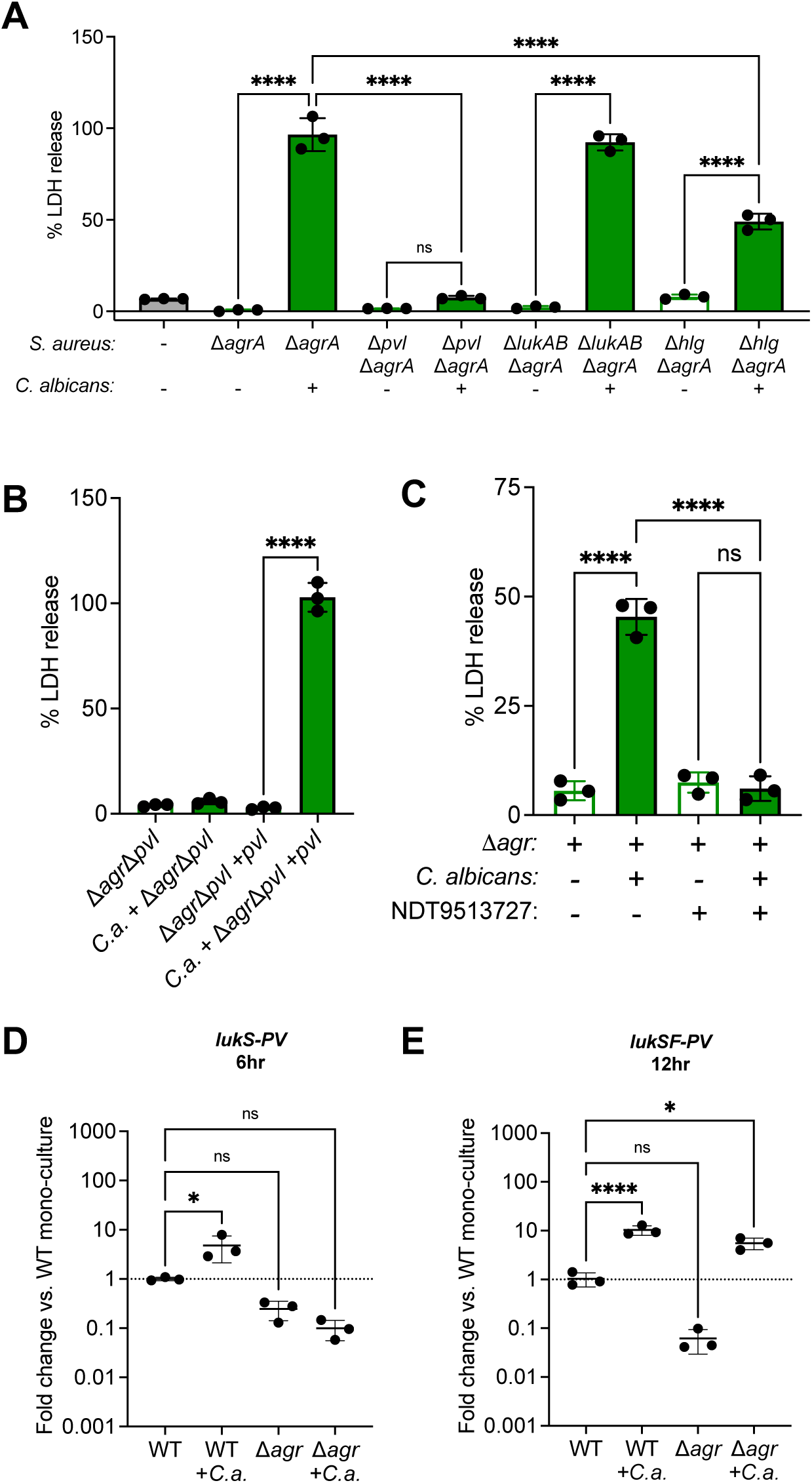
PVL is required for *S. aureus* Δ*agr* cytotoxicity induced by *C. albicans*. (A-B) Primary human CD14^+^ monocytes were exposed to supernatants from *S. aureus* strains cultured with and without *C. albicans*. % LDH release was quantified at 24 hours after supernatant exposure and calculated by standardizing LDH values in each well against cells lysed with 10% Triton-X 100 as a positive cell death control. (C) Monocytes were treated with 10 μM NDT9513727 or DMSO prior to the addition of culture supernatant and % LDH release was quantified at 6 hours. (D-E) qRT-PCR analysis of *lukS-PV* transcript levels for cultures at 6 hours (D) and 12 hours (E) of growth. *S. aureus* WT = wild-type; Δ*agr* = *agrACBD::tet;* Δ*agrA = agrA::bursa*; Δ*pvl = lukSF-PV::spc*; Δ*AB* = Δ*lukAB*; Δ*hlg = hlgACB::tet,* Δ*agr*Δ*pvl* +*pvl* = *agrACBD::tet pvl::spc attC*::pJC1111-P*pvl.* Values are plotted as fold change relative to the WT mono-culture samples. For bar graphs, ****p<0.001 by one-way ANOVA with Tukey’s multiple comparisons test; for qRT-PCR, **p<0.01, ***p<0.005, and ns (not significant) by one-way ANOVA with Dunnett’s multiple comparisons test. # represents p<0.001 compared to *S. aureus* Δ*agrA* co-cultured with *C. albicans.* N = 3 wells or replicates per condition, bars = mean±SD.

To confirm the role of PVL in driving *S. aureus* Δ*agr* co-culture cytotoxicity, we utilized an *agrBDCA::tet*/*lukSF-PV::spc* mutant (designated Δ*agr*Δ*pvl*) to evaluate cytotoxicity of culture supernatants towards human monocytes at 24 hours after supernatant exposure. *C. albicans* does not induce Δ*agr*Δ*pvl* cytotoxicity towards human monocytes (**Fig. 4B**). Complementation of *lukSF-PV* at a neutral site on the chromosome restores the ability of *C. albicans* co-culture to induce cytotoxicity of *S. aureus* Δ*agr*Δ*pvl*, again highlighting the critical role for PVL in driving greater *C. albicans-*induced human monocyte cytotoxicity (**Fig. 4B**).

PVL is a bi-component pore-forming toxin that consists of two subunits, LukS-PV and LukF-PV. LukS-PV selectively binds the human isoform of the C5aR1 receptor on host cells to initiate pore-formation (37). Previous work reveals that pharmacological blockade of C5aR1 on human neutrophils prior is sufficient to inhibit PVL-mediated cell lysis (52, 53). Because we observed that PVL is required for *S. aureus* Δ*agr* co-culture cytotoxicity, we hypothesized that human cell death is dependent on PVL binding to the C5aR1 receptor. To test this, human monocytes were pre-treated with NDT9513727, a C5aR1 negative allosteric modulator (53, 54), prior to supernatant exposure. Although *C. albicans* induces significant *S. aureus* Δ*agr* cytotoxicity towards mock-treated monocytes, pre-treatment of monocytes with NDT9513727 was sufficient to inhibit cell death (**Fig. 4C**).

We next evaluated PVL transcript levels in *S. aureus* wild-type and Δ*agr* cultures by qRT-PCR. *C. albicans* co-culture enhances *lukS-PV* transcript levels in *S. aureus* wild-type at both 6 and 12 hours of culture growth (**Fig. 4D-E**). *S. aureus* Δ*agr* mono-culture has reduced *lukS-PV* transcript compared to *S. aureus* wild-type, and *C. albicans* co-culture significantly enhances transcript levels at 12 hours of co-culture (**Fig. 4D-E**). Taken together, our data reveal that *C. albicans* enhances *lukSF-PV* expression in *S. aureus* Δ*agr,* and PVL is required for *S. aureus* Δ*agr* co-culture cytotoxicity towards human monocytes.

### Mutation of the SaeR binding site upstream of *pvl* reduces human cell cytotoxicity induced by co-culture

We determined that both *S. aureus* SaeRS and the human-selective toxin PVL are required for Agr-independent human cell death induced by *C. albicans*. Activation of the SaeRS system results in increased phosphorylation of SaeR, which drives greater transcription of target genes (55). Therefore, we sought to determine if SaeR regulation of PVL contributes to cytotoxicity induced by *C. albicans*. To test this, we first identified and mutated the SaeR binding site consensus sequence upstream of the *pvl* locus in *S. aureus* Δ*agr,* generating the strain Δ*agr/pvl^sbm^.* We cultured *S. aureus* Δ*agr* and Δ*agr/pvl^sbm^* with and without *C. albicans* and tested cytotoxicity towards human monocytes. While *C. albicans* induces some cytotoxicity of *S. aureus* Δ*agr/pvl^sbm^*, the amount of cell death is significantly reduced compared to cytotoxicity induced by *S. aureus* Δ*agr* co-culture (**Fig. 5**). To complement the *S. aureus* Δ*agr/pvl^sbm^* strain, we reverted the mutated SaeR binding site sequence back to wild-type, generating *S. aureus* Δ*agr/pvl^rev^*. Co-culture with *S. aureus* Δ*agr/pvl^rev^*induces significantly higher levels of human cell death, as compared to *S. aureus* Δ*agr/pvl^sbm^* co-culture (**Fig. 5**). These data suggest that *C. albicans* enhances PVL-mediated human cell death in part through direct SaeR regulation at the *pvl* locus.

**Figure 5.**
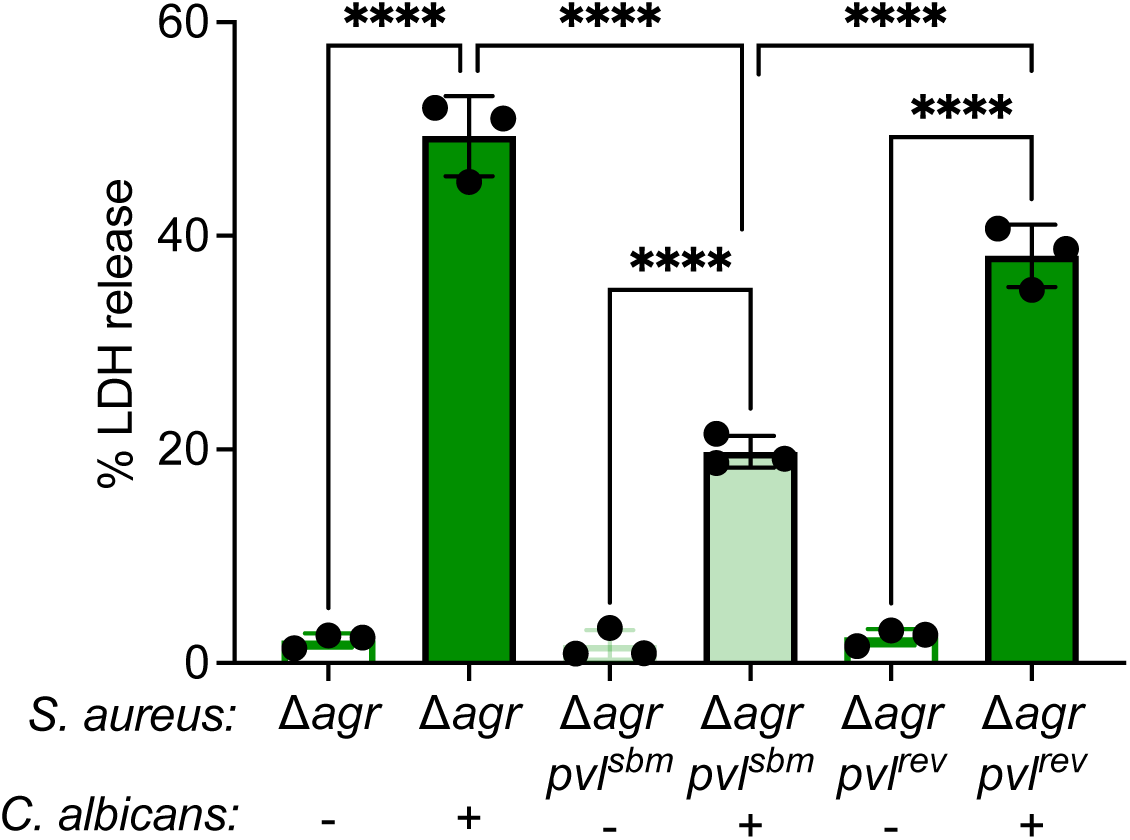
Mutation of the SaeR binding site upstream of *pvl* reduces *S. aureus* Δ*agr* cytotoxicity. Primary human CD14^+^ monocytes were exposed to supernatants from *S. aureus* Δ*agr,* Δ*agr/pvl^sbm^*(SaeR binding site mutant strain), and Δ*agr/pvl^rev^* (SaeR binding site wild-type revertant strain) cultured with and without *C. albicans*. % LDH release was quantified at 24 hours after supernatant exposure and calculated by standardizing LDH values in each in each well against cells lysed with 10% Triton-X 100 as a positive cell death control. *S. aureus* Δ*agr* = *agrACBD::tet; pvl^sbm^ =* SaeR binding site mutant upstream of *pvl*; *pvl^rev^ =* revertant strain. ****p<0.001, and ns (not significant) by one-way ANOVA with Tukey’s multiple comparisons test. N = 3 wells or replicates per condition, bars = mean±SD.

### *C. albicans*-enhanced cytotoxicity is conserved across multiple *S. aureus* clinical isolates

To determine if enhanced cytotoxicity following co-culture is conserved among *S. aureus* strains, we tested multiple *S. aureus* clinical isolates. We chose nine *S. aureus* isolates that were classified as USA300 sequence type 8, which is the same background as the LAC* wild-type strain we use in this study (56). Primary human monocytes were exposed to mono– and co-culture supernatants, and LDH release was evaluated at 6 hours. *C. albicans* significantly enhances cytotoxicity towards human monocytes for all of the strains tested except for isolate VUSA11, which induces high levels of cell death for both mono– and co-culture (**Fig. 6A**). By 24 hours after supernatant exposure, all *S. aureus* wild-type isolates cultured alone or with *C. albicans* induce maximal cell death (**Fig. S7**).

**Figure 6.**
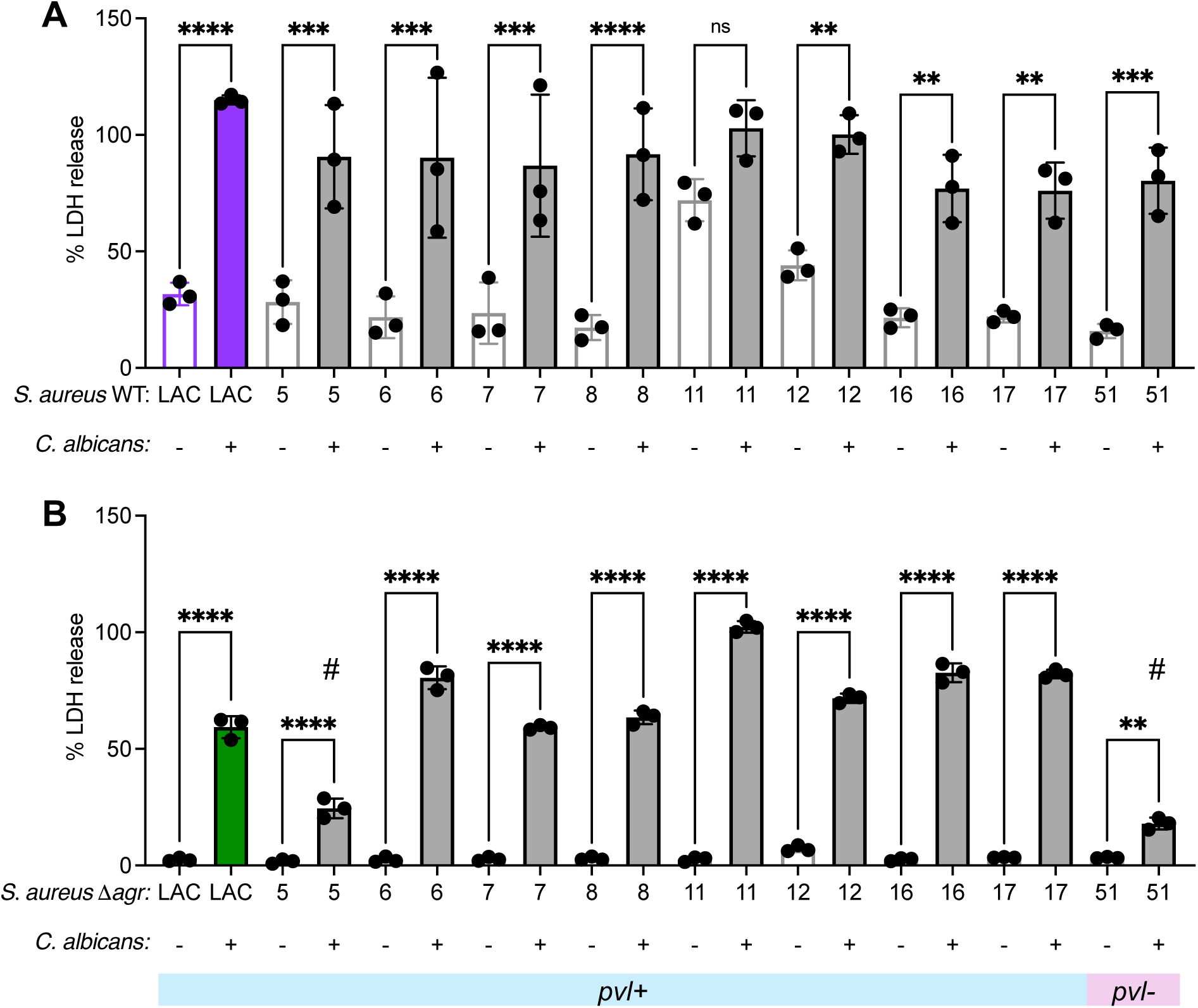
*C. albicans* co-culture enhances cytotoxicity of *S. aureus* USA300 clinical isolates and Δ*agr* strains. Primary human CD14^+^ monocytes were exposed to supernatants from (A) *S. aureus* LAC* and USA300 clinical isolates and (B) the isogenic Δ*agr* mutant for each isolate, both cultured with and without *C. albicans*. % LDH release was quantified at (A) 6 hours or (B) 24 hours after supernatant exposure and calculated by standardizing LDH values in each in each well against cells lysed with 10% Triton-X 100 as a positive cell death control. All strains are *pvl+* (encode *lukSF-PV*) except for clinical isolate 51. **p<0.01, ****p<0.001, and ns (not significant) by one-way ANOVA with Tukey’s multiple comparisons test. # represents p<0.001 compared to *S. aureus* LAC* Δ*agr* co-cultured with *C. albicans.* N = 3 wells or replicates per condition, bars = mean±SD.

To evaluate if *C. albicans* induces cytotoxicity of *S. aureus* isolates with *agr* mutation, we generated Δ*agr* (*agrBDCA::tet*) strains for each of the clinical isolates in **Fig. 6A**. Human monocytes were exposed to mono– and co-culture supernatants for each Δ*agr* isolate, and LDH release was measured at 24 hours. Mutation of the Agr system ablates cytotoxicity towards human monocytes for each of the *S. aureus* clinical isolates when grown in mono-culture (**Fig. 6B**). *C. albicans* induces cytotoxicity for seven of the nine clinical isolates to the same extent or greater than LAC* Δ*agr* co-culture. Two clinical isolates (VUSA5 and VUSA51) have significantly lower co-culture cytotoxicity compared to LAC* Δ*agr* co-culture. We next performed whole genome sequencing for all *S. aureus* isolates to try and identify why the VUSA5 and VUSA51 were less cytotoxic following co-culture. We determined that VUSA5 was PVL negative and VUSA51 had a mutation in *arlS.* Taken together, these data suggest that *C. albicans* enhances cytotoxicity of multiple USA300 clinical isolates and the isogenic Δ*agr* strains.

### *C. albicans* clinical isolates broadly enhance *S. aureus* wild-type and Δ*agr* cytotoxicity

High levels of phenotypic variation exist among *C. albicans* clinical isolates. Therefore, we examined the ability of *C. albicans* isolates to enhance *S. aureus* wild-type and Δ*agr* cytotoxicity compared to *C. albicans* SC5314, the wild-type reference strain used throughout this study. *S. aureus* wild-type and Δ*agr* were cultured with *C. albicans* SC5314 or *C. albicans* clinical isolates. Because *C. albicans* co-culture can alter media pH and impact *S. aureus* virulence regulation (57), we first measured the pH of the co-cultures and confirmed that the *C. albicans* clinical isolates neutralize the media to a similar extent as *C. albicans* SC5314 (**Fig. S8**). Next, we observed that each *C. albicans* isolate tested enhances cytotoxicity of *S. aureus* wild-type towards human monocytes, as determined by LDH release 6 hours after supernatant exposure (**Fig. 7A**). Two of the *C. albicans* clinical isolates (Ca91 and Ca97) enhance *S. aureus* wild-type cytotoxicity to a greater extent compared to the cytotoxicity observed following co-culture with *C. albicans* SC5314. The other eleven isolates enhance *S. aureus* wild-type cytotoxicity to the same level as *C. albicans* SC5314. We also tested cytotoxicity of the same *C. albicans* clinical isolates co-cultured with *S. aureus* Δ*agr*. We observed that each clinical isolate induces cytotoxicity of *S. aureus* Δ*agr* during co-culture (**Fig. 7B**). However, nine of the thirteen isolates tested induce significantly less cytotoxicity relative to *C. albicans* SC5314 co-culture. These data indicate that the ability to enhance *S. aureus* cytotoxicity is broadly conserved among *C. albicans* isolates, but the magnitude that *C. albicans* isolates induce Δ*agr* cytotoxicity is more variable.

**Figure 7.**
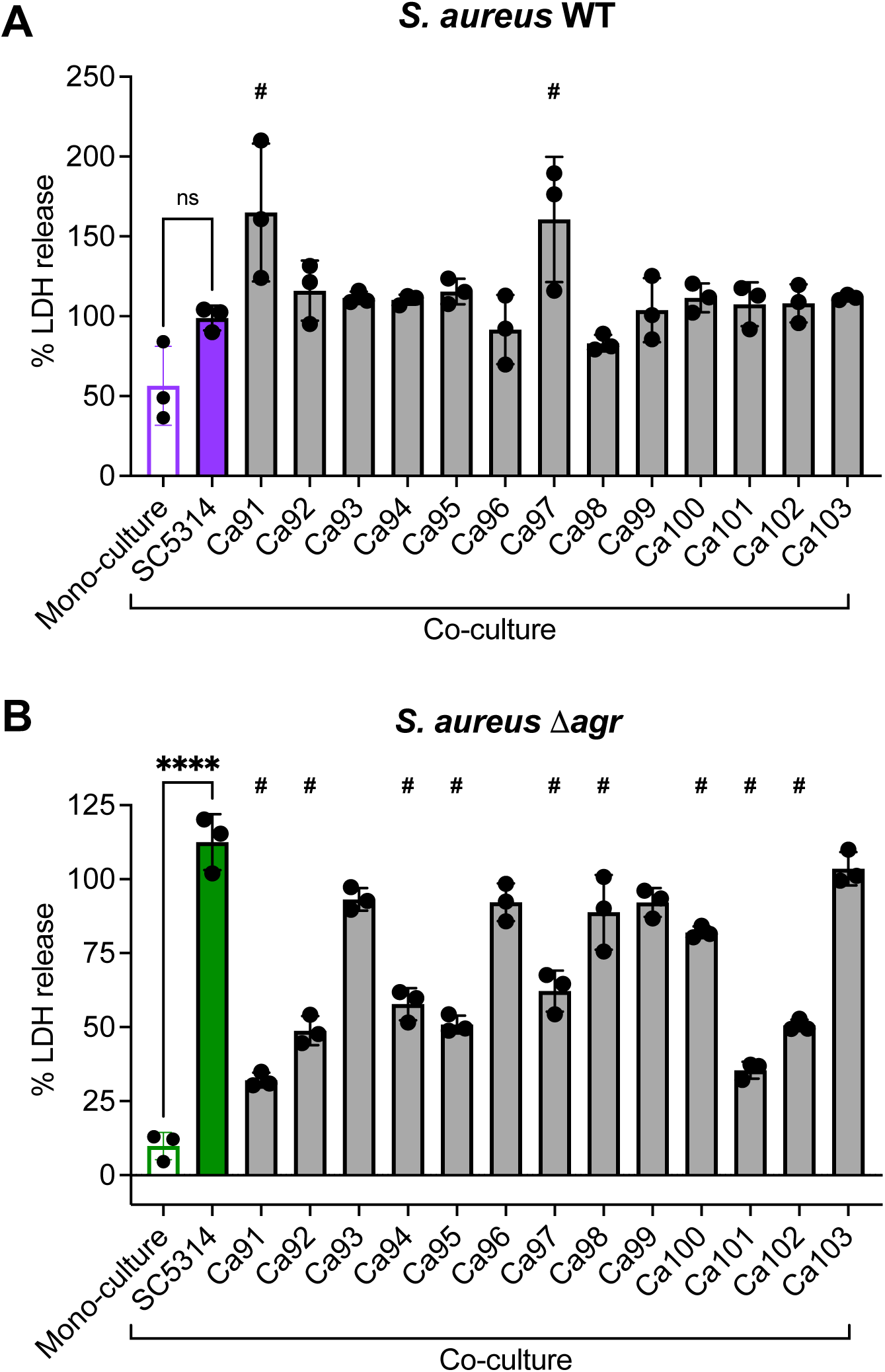
*C. albicans* clinical isolates broadly enhance *S. aureus* wild-type cytotoxicity but variably induce *S. aureus* Δ*agr* cytotoxicity. Primary human CD14^+^ monocytes were exposed to supernatants from (A) *S. aureus* wild-type or (B) *S. aureus* Δ*agr* that was cultured with and without *C. albicans* SC5314 or the indicated *C. albicans* clinical isolate. % LDH release was quantified at (A) 6 hours or (B) 24 hours after supernatant exposure and calculated by standardizing LDH values in each in each well against cells lysed with 10% Triton-X 100 as a positive cell death control. ****p<0.001 and ns (not significant) by one-way ANOVA with Tukey’s multiple comparisons test. # represents p<0.0001 compared to *S. aureus* co-cultured with *C. albicans* SC5314. N = 3 wells or replicates per condition, bars = mean±SD.

## Discussion

Using both human and murine cells, we determined that *C. albicans* enhances *S. aureus* virulence by engaging two staphylococcal virulence regulatory systems to induce cell death with host species-specific effects. While *C. albicans* enhances *S. aureus* cytotoxicity towards murine monocytes through the activity of Agr-regulated toxins, we uncovered that *C. albicans* unexpectedly induces cytotoxicity of the typically non-toxic *S. aureus agr* mutant, with selectivity towards human monocytes and neutrophils. Our data reveal that *C. albicans* can activate the *S. aureus* SaeRS two-component system independently of Agr, and SaeRS activation is required for induced cytotoxicity towards human cells. We further identified the SaeRS-regulated toxin PVL as the main factor contributing to human monocyte cell death when *S. aureus* Δ*agr* is co-cultured with *C. albicans.* Co-culture induces significant cytotoxicity for multiple *S. aureus* clinical isolates with and without *agr* mutation. Additionally, multiple *C. albicans* clinical isolates are able to enhance cytotoxicity of *S. aureus* wild-type and Δ*agr* during co-culture.

The *S. aureus* Agr system is regarded a major virulence regulatory system due in part to the large repertoire of secreted toxins whose production is increased following Agr activation (33, 36). Previous work revealed that *C. albicans* enhances *S. aureus* Agr activation to elaborate α-hemolysin production, contributing to greater mortality in a murine model of intraabdominal co-infection (31). Our results expand on this finding by demonstrating that *C. albicans* co-culture enhances *S. aureus* cytotoxicity towards murine monocytes via an Agr-dependent mechanism requiring the α-PSM cytolytic peptides and the leukocidin γ-hemolysin. We did not observe a significant role for α-hemolysin in mediating enhanced cytotoxicity towards monocytes, but this toxin likely plays a role in regulating enhanced cytotoxicity towards other murine cell types, especially during intra-abdominal co-infection (31). We also discovered that *C. albicans* enhances *S. aureus* virulence independently of the Agr system via a mechanism requiring SaeRS. SaeRS is a two-component system that regulates over 20 virulence factors in *S. aureus* (58). Because inactivation of Agr reduces Sae transcript levels, the SaeRS two-component system was initially thought to be activated by Agr (59). Supporting this framework for regulation, we observe reduced *saeP* transcript levels in our *S. aureus* Δ*agr* strain compared to wild-type at both 6 and 12 hours of culture. This may also explain our observation that while *C. albicans* co-culture significantly enhances *S. aureus* SaeRS reporter activation in *S. aureus* Δ*agr* relative to mono-culture, the overall activation is about 3-fold lower compared to levels for *S. aureus* wild-type co-cultured with *C. albicans.* However, it is important to note that SaeRS transcript levels do not always correlate with the level of SaeR activity (60). While we observe an Agr-dependent effect on *saeP* transcript levels, our data also support a significant role for SaeRS in driving human monocyte cytotoxicity independently of *S. aureus* Agr following *C. albicans* co-culture. Other studies have also demonstrated a role for SaeRS in promoting *S. aureus* virulence independently of the Agr system (45, 50). Thus, while some SaeRS activity may be regulated downstream of Agr, SaeRS can also be activated by *C. albicans* independently of the Agr system.

We further determined that the SaeRS-regulated leukocidin PVL was required for the enhanced cytotoxicity of *S. aureus* Δ*agr* following *C. albicans* co-culture. *C. albicans* failed to induce cytotoxicity of *S. aureus* Δ*agr/*Δ*pvl* towards human monocytes, and cytotoxicity could be restored through complementation of *lukSF-PV*. We also found that pre-treatment of primary human monocytes with NDT9513727, a negative allosteric modulator used as a potent C5aR1 inhibitor, was sufficient to inhibit *S. aureus* Δ*agr* co-culture cytotoxicity. PVL is species-specific, with high levels of activity towards human and rabbit cells, but poor activity towards murine cells (37). Cell and species specificity is mediated through binding of the toxin S-subunit to the human isoform of C5aR1 and the F-subunit to the human isoform of CD45 (37, 52). Because of the human-specific activity of PVL, traditional murine infection models would not be an effective system for testing the role of enhanced PVL production during *C. albicans-S. aureus* co-infection. Transgenic mice that express the human isoform of the C5aR1 receptor have been employed to evaluate *S. aureus* pathogenesis for human-selective toxins, but PVL has a limited impact in this model due to the lack of the human CD45 co-receptor (52). Recently the huBRGSF humanized mouse strain (BALB/c Rag2^-/-^Il2rg^-/-^Sirpa^NOD^Flk2^-/-^) was used to test the role of PVL in the pathogenesis of *S. aureus* osteomyelitis (61). These mice could also be used to evaluate how interactions with *C. albicans* impact *S. aureus* Δ*agr* pathogenesis during infection.

Our data suggest a model in which *C. albicans* induces activation of SaeRS in both *S. aureus* wild-type and Δ*agr* during co-culture, and enhanced SaeRS activation in Δ*agr* induces greater PVL production that drives human monocyte cell death. Mutation of the SaeR binding site upstream of the *pvl* locus successfully reduced cytotoxicity driven by co-culture, supporting this model. However, *C. albicans* still induces some cytotoxicity of *S. aureus* Δ*agr/pvl^sbm^.* This could be due to SaeR partially binding the mutated sequence with reduced affinity, or there may be additional, indirect SaeR regulation of *pvl* expression. SaeRS regulates multiple toxins, including several with selective activity towards human cells, but we only identified a role for PVL in regulating enhanced Agr-independent cytotoxicity (49). If *C. albicans* activates SaeRS, then there should be enhanced production of other toxins that induce human monocyte cell death, such as γ-hemolysin or LukAB, and not just PVL. One possibility for the sole involvement of PVL in our model is that our primary human monocyte culture methods may result in higher levels of C5aR1 (the PVL receptor) displayed on the surface of the monocytes as compared to the receptors for other toxins, leading to greater cytolytic effects driven by PVL. This may be occurring because we culture our monocytes in macrophage colony stimulating factor (M-CSF) for six days prior to evaluating cytotoxicity, and M-CSF induces higher levels of C5aR1 on the surface of macrophages (62). Another possibility is that *C. albicans* induces additional transcriptional or post-transcriptional changes in *S. aureus* Δ*agr* that result in selectively greater production of PVL, as opposed to other SaeRS-regulated human specific toxins. Finally, it is also possible that because activation of SaeRS is lower in Δ*agr* co-culture compared to wild-type co-culture, other SaeRS-regulated toxins may be present in culture supernatant but at sub-lytic levels. A limitation to our study is that we only evaluated the species-specific effect of secreted toxins by applying filter-sterilized culture supernatants to host cells. We chose this reductionist approach to evaluate impacts of *C. albicans* co-culture on the *S. aureus* secreted toxin repertoire. However, additional species-specific effects of *C. albicans-S. aureus* interactions may be revealed through *in vitro* and *in vivo* infections with live organisms, including revealing a potential role for other SaeRS-regulated toxins.

The presence of *lukSF-PV,* the locus that encodes the gene for PVL, is commonly associated with *S. aureus* isolates from the USA300 lineage, which is the background for the strain used in our study (56). To test if *C. albicans* broadly enhances cytotoxicity of USA300 strains with and without Agr, we included nine clinical isolates that were the same sequence type and clonal complex as our laboratory USA300 strain. *C. albicans* co-culture significantly enhanced cytotoxicity of eight of the nine isolates towards human monocytes, while the ninth isolate was already highly cytotoxic in mono-culture. Furthermore, mutation of the *agr* locus rendered each isolate non-toxic in mono-culture, and *C. albicans* enhanced cytotoxicity of seven *agr* mutant isolates to the same level as the LAC* *agr* mutant. Two of the *agr* mutant strains (VUSA5 and VUSA51) had significantly lower cytotoxicity as compared to the LAC* strain following co-culture with *C. albicans.* Of note, VUSA51 is PVL– and VUSA5 is PVL+, but VUSA5 harbors a point mutation in *arlS,* which introduces an early stop codon. ArlRS is a two-component system that has been shown to regulate PVL expression (63). Therefore, it is possible that VUSA5 is unable to fully produce PVL due to the mutation in *arlS*. These results highlight the importance of PVL in driving enhanced virulence of *S. aureus agr* mutant strains towards human monocytes and demonstrate the ability of *C. albicans* to enhance the virulence of *S. aureus agr* null isolates. Strains with Agr dysfunction have been isolated from chronic infections, including from patients with cystic fibrosis or bacteremia, and so interactions with *C. albicans* during these infections could promote virulence of *agr* null strains.

The mechanism by which *C. albicans* induces *S. aureus* SaeRS activation during co-culture is unknown. Recent studies revealed that *C. albicans* enhances *S. aureus* Agr activation through modifications to the extracellular environment. *C. albicans* maintains a neutral pH of the co-culture media through the release of ammonia as a byproduct of amino acid metabolism, and *S. aureus* Agr activation is maximized at neutral pH (57, 64). Additionally, *C. albicans* can induce activation of Agr during co-culture by metabolizing and depleting ribose from the co-culture environment which alleviates a ribose-repressive effect on the Agr system (65). Several factors have been shown to activate SaeRS in *S. aureus* grown alone, although the precise mechanism of action is incompletely understood. SaeRS transcriptional activity was previously shown to be higher when *S. aureus* is grown in neutral pH compared to acidic pH (66). However, we identified several *C. albicans* clinical isolates that induce significantly less Δ*agr* cytotoxicity compared to *C. albicans* SC5314, but these strains are still able to maintain neutral pH during co-culture. This suggests that neutral pH alone is not sufficient to induce SaeRS activation during co-culture. Additionally, each *C. albicans* isolate tested was able to enhance cytotoxicity of *S. aureus* wild-type during co-culture. Therefore, *C. albicans* induction of SaeRS may be occurring via a different mechanism than *C. albicans* enhancement of Agr activation.

Overall, this study reveals that interactions with the common co-colonizing and co-infecting fungus *C. albicans* lead to activation of two major virulence regulatory systems in *S. aureus,* Agr and SaeRS. SaeRS was required for cytotoxicity towards human monocytes, demonstrating species-specific effects of *S. aureus-C. albicans* interactions. Because we revealed that *C. albicans* activates SaeRS in *S. aureus* Δ*agr* mutants, *agr-*null isolates may also be susceptible to virulence regulation by *C. albicans* in the context of human infections. This study underscores the importance of evaluating how *S. aureus* interactions with *C. albicans* impact virulence towards human cells in addition to murine infection models for a more complete and translationally impactful view of how polymicrobial interactions can shape *S. aureus* disease.

## Materials and Methods

### Ethics statement

All experiments involving animals for cell isolation were reviewed and approved by the Institutional Animal Care and Use Committee of Vanderbilt University Medical Center and performed according to NIH guidelines, the Animal Welfare Act, and US Federal Law. Human primary peripheral blood mononuclear cells were commercially sourced from Zen-Bio (a BioIVT company), a commercial vendor that de-identifies their products so donor personal information is not publicly available. Primary human neutrophils were isolated from peripheral whole blood obtained from consenting healthy donors by the Vanderbilt Vaccine Research Program, according to the IRB protocol 070258. All *C. albicans* and *S. aureus* clinical isolates were received in a deidentified manner to protect personal health information.

### Microbial strains and culture conditions

Strain information for *S. aureus* can be found in Table S1 and *C. albicans* in Table S2. Experiments were performed with *S. aureus* USA300 lineage strain LAC* (AH1263), an erythromycin– and tetracycline-sensitive derivative of LAC, as the wild-type (WT) strain. All *S. aureus* strains used in this study are in this strain background, except for the clinical isolate strains, which were obtained in a de-identified fashion from a separate IRB-approved study. Mutants generated in this study were created using phi-85-mediated phage transduction. Specifically, *agrA::tn* (Δ*agrA*) from the NARSA strain collection was transduced into Δ*lukAB*, *pvl::spc* (Δ*pvl*), and *hlgACB::tet* (Δ*hlg*) strains. Additionally, *agrACDB::tet* (Δ*agr*) was transduced into the P*_saeP_*-*gfp* (*attC*::P*_sarA_*-*sod*RBS*-mCherry* SAUSA300_RS05730::P*_saeP_*-*gfp*) and P*_blank_*-*gfp* (*attC*::P_sarA_-*sod*RBS*-mCherry* SAUSA300_RS05730::*gfp*) strains. *S. aureus* Δ*pvl* and each of the *S. aureus* clinical isolates were also transduced with *agrACDB::tet* (Δ*agr*) to generate Δ*agr/*Δ*pvl* and clinical isolate Δ*agr* strains, respectively. Complementation of Δ*agr/*Δ*pvl* was done using pJC1111-p*pvl,* which was created in (67) by cloning the genes encoding *pvl* and their native promoter region into pJC1111 and integrating this construct into the SaPI1 *attC* site in RN4220. We transduced this complementation construct(67) into *S. aureus* Δ*agr/*Δ*pvl,* generating Δ*agr/*Δ*pvl +pvl* (*agrACBD::tet lukSF-PV::spc* SaPI1 *attC::*p*pvl-pvl*). *C. albicans* strain SC5314 was used in all experiments as the wild-type reference strain. *C. albicans* clinical isolates were obtained from the *micro*VU biorepository at Vanderbilt University in a de-identified fashion. *S. aureus* strains were grown on tryptic soy agar (TSA) plates before inoculation into 5 mL tryptic soy broth (TSB) in 15 mL conical tubes, which were grown overnight with shaking at 180 rpm and incubation at 37°C. Erythromycin (10 μg/mL), tetracycline (2 μg/mL), spectinomycin (100 μg/mL), or cadmium chloride (0.1 mM) were added to cultures for strains that possess the corresponding resistance cassettes. *C. albicans* strains were grown first on yeast peptone dextrose (YPD) agar plates and then inoculated in 5 mL YPD broth in 50 mL Erlenmeyer flasks with shaking at 200 rpm and incubation at 30°C overnight.

### Generating *S. aureus* SaeR site binding mutant and revertant strains

Construction of a strain harboring a SaeR-binding site mutation upstream of *pvl* and the associated revertant strain were created as described previously (68). Briefly, the SaeR binding motif sequence upstream of *pvl*, GTTAA(N_6_)TTAA, was mutated to GGCGG(N_6_)GGAG by overlapping PCR using primers pvl1F (5’-ATGGAATTCGAATCCGCCAGTGCCAGC-3’), pvl2R (5’-CATTTCCTTTCTTTATAAATTTTATTACATTTTTATACTCCACCTTTCCGCCTTTTAATAA AATTAA-3’), pvl3F (5’-TAATTTTATTAAAAGGCGGAAAGGTGGAGTATAAAAATGTAATAAAATTTATAAAGAA-3’), and pvl4R (5’-ATGGGTACCGAAGGATTGAAACCACTGTGTAC-3’) and *S. aureus* wild-type genomic DNA as the template. The mutated fragment was cloned into pKOR1 vector at the EcoRI and KpnI restriction sites, and allelic replacement on the chromosome of *S. aureus* wild-type was performed as described previously (69), generating the binding site mutant (designated *pvl^sbm^*). To generate the revertant strain, a wild-type fragment was amplified from *S. aureus* wild-type via PCR using primers pvl1F and pvl4R. The wild-type fragment was cloned into pKOR1 at the EcoRI and KpnI restriction sites. Allelic replacement was performed in the *pvl^sbm^* strain, generating the revertant strain (designated *pvl^rev^*). Successful mutation of the SaeR binding site and reversion back to wild-type was confirmed by PCR and sequencing. Following confirmation, *S. aureus pvl^sbm^* and *pvl^rev^* were transduced with *agrACDB::tet* (Δ*agr*) using phi-85 phage, generating *S. aureus* Δ*agr/pvl^sbm^* and Δ*agr/pvl^rev^*.

### Mono– and co-culture supernatant preparation

1 mL of overnight cultures of *S. aureus* and *C. albicans* were pelleted, and pellets were washed once in 1 mL 1X phosphate buffered saline (PBS). Cells were pelleted again, and *S. aureus* cells were resuspended in 1 mL PBS and *C. albicans* cells were resuspended in 500 μL PBS. To set up co-cultures, 50 μL of *S. aureus* and 50 μL of *C. albicans* cells were added to 5 mL RPMI-g, which is RPMI1640 (Gibco) supplemented with 1% casamino acids and 2% D-glucose, in a 50 mL conical tube. To set up mono-cultures, 50 μL of either *S. aureus* or *C. albicans* and 50 μL of RPMI-g media were added to 5 mL RPMI-g in a 50 mL conical tube. Cultures were grown for 15 hours at 37°C with shaking at 180 rpm. A portion of culture was removed before and after 15 hours and plated on selective media to determine microbial CFU. For growth curves, a portion of culture was removed and plated every 3 hours over 15 hours of growth. After 15 hours growth, cultures were centrifuged at 4000 rpm for 10 minutes to pellet microorganisms. Cultures were kept on ice while each supernatant was filter-sterilized through a 0.22 μm filter into a new tube. Filter-sterilization was repeated, and a portion of the culture supernatant was plated to confirm there was no microbial growth. Supernatants were aliquoted and used either immediately or stored at –80°C until use in cytotoxicity assays.

### Murine bone marrow-derived monocyte isolation and culture

Murine bone marrow-derived monocytes (BMDM) were isolated from wild-type C57BL6/J female mice aged 8-12 weeks. Whole bone marrow (WBM) was flushed from femurs using 5 mL MEM-α medium (Gibco). WBM cells were pelleted by centrifugation at 1500 rpm for 5 minutes. Media was decanted and pellets were resuspended in 1 mL ACK buffer (KD Biomedicals) and incubated at room temperature for 5 minutes to allow for erythrocyte lysis. After incubation, 9 mL of 1X PBS was added and WBM was filtered through a 70 μm nylon filter into a new conical tube. Cells were pelleted by centrifugation at 1500 rpm for 5 minutes. Media was decanted and WBM was resuspended to a concentration of 8 x 10^6^ cells/mL in complete MEM-α medium, which is MEM-α supplemented with 10% fetal bovine serum and 1X penicillin/streptomycin cocktail (Corning). In 10 cm tissue culture-treated dishes, 1 mL of cells were combined with 8 mL of complete media and 2 mL of supernatant derived from the CMG 14-12 cell line as a source of M-CSF. WBM cells were allowed to differentiate for 4 days with incubation at 37°C and 5% CO_2_, after which time cells were removed by scraping. BMDM were resuspended in FBS and stored in liquid nitrogen. Prior to cytotoxicity assays, frozen BMDM were rapidly thawed at 37°C and washed in complete MEM-α medium. Cells were seeded in 96-well tissue culture plates at a concentration of 50,000 cells in 200 μL per well. Cells were cultured an additional two days prior to use in cytotoxicity assays.

### Negative selection for human CD14^+^ monocytes

Fresh peripheral blood mononuclear cells (PBMC) isolated from healthy human donors were received from ZenBio. Cells were either used upon arrival for negative selection or aliquoted and stored frozen in CryoStor cell preservation media (Sigma). If starting the negative selection from frozen cells, cells were thawed at 37°C and immediately transferred to complete MEM-α media. Negative selection for CD14^+^ monocytes was performed using the Classical Monocyte Isolation Kit, Human (Miltenyi) and following manufacturer instructions. Briefly, cells were pelleted by centrifugation at 300 x *g* for 10 minutes and then resuspended in 30 μL of buffer (1X PBS + 20% FBS + 0.5M EDTA) for every 10^7^ cells. 10 μL each of Fc block and 10 μL of antibody cocktail were added for every 10^7^ cells, followed by 5 μL of thrombocyte removal agent added for every 10^7^ cells. After 5 minutes incubation at room temperature, 30 μL of buffer and 20 μL of anti-biotin microbeads were added for every 10^7^ cells. Cells were mixed gently and incubated at room temperature for 5 min. Negative selection was performed with the LS column (Miltenyi), where the CD14^+^ cell portion is contained within the flow-through. The column was washed with 3 mL buffer, and the CD14^+^ cells were centrifuged 300 x *g* for 10 minutes. Cells were resuspended in complete MEM-α media supplemented with 50 ng/mL of recombinant human M-CSF (R&D Systems). For cytotoxicity assays, cells were resuspended at a concentration of 2.5 x 10^5^ cells/mL and 96-well tissue culture plates were seeded with 5.0 x10^4^ cells in 200 μL per well. For SYTOX imaging assays, cells were resuspended at a concentration of 5×10^5^ cells/mL, and μ-Slide 8 Wells (Ibidi) were seeded with 1.5×10^5^ cells in 300 μL per chamber. Cells incubated at 37°C with 5% CO_2_ for six total days, with a complete media change at three days after seeding. Cells were used in cytotoxicity assays or SYTOX assays six days after seeding.

### Human neutrophil isolation

Primary neutrophils were isolated from peripheral blood of healthy human donors as previously described (70). Briefly, 30 mL of peripheral blood was collected into EDTA-containing vacuum tubes by standard venipuncture. Blood was mixed with 20 mL of a 3% dextran-0.9% sodium chloride solution and incubated at room temperature for 20 minutes to allow erythrocyte sedimentation. The top layer was removed and centrifuged at 250 x g for 10 minutes. The cell pellet was resuspended in 25 mL 0.9% sodium chloride and underlaid with 10 mL Ficoll-Paque Plus (GE Healthcare). Centrifugation at 400 x g for 40 minutes at 20C was performed to pellet neutrophils and remaining erythrocytes. All other layers were aspirated and the pellet was resuspended in 20 mL of 0.2% sodium chloride for red blood cell lysis. After 30 seconds, 20 mL of 1.6% sodium chloride was added to restore tonicity. Cells were pelleted at 250 x *g* for 6 minutes, and the erythrocyte lysis step was repeated once more. Neutrophils were resuspended in RPMI1640 (Gibco) at a final concentration of 5 x 10^5^ cells/mL. 96-well tissue culture plates were seeded at a concentration of 100,000 cells in 200 μL per well. Cells were used immediately in the cytotoxicity assay.

### Culture of cell lines

A549 cells were obtained from the American Type Culture Collection (ATCC) and primary human lung microvascular endothelial cells (HLMVEC) were obtained from Promocell. All cells were propagated according to manufacturer recommendations. A549 were cultured in Dulbecco’s MEM (DMEM) supplemented with 10% FBS and 1X penicillin/streptomycin, and HLMVEC were cultured in Endothelial Cell Growth Medium MV2 (Promocell). Media was exchanged every 2-3 days for both cell types. Two days prior to cytotoxicity assays, A549 cells were seeded in 96 well tissue culture plates at a density of 15,000 cells in 200 μL per well. Prior to use in cytotoxicity assays, HLMVEC were seeded in 96 well plates coated with Attachment Factor, and cells were incubated for another 2-3 days to allow for the formation of tight junctions. All cells were grown or maintained at 37°C and 5% CO_2_.

### Cytotoxicity assay

Culture medium was removed from each well of A549, HLMVEC, BMDM, CD14^+^, or neutrophil cells that were isolated and seeded as described above. Fresh media for each cell type (without added FBS or penicillin/streptomycin) was added back to each well at a volume of 150 μL or 160 μL, for testing supernatants at a 25% (A549 cells) or 20% (all other cell types) volume/volume concentration. MEM-α for BMDM and CD14^+^ were supplemented with an M-CSF source. Where indicated, CD14^+^ monocytes were treated with 10 μM NDT9513727 (R&D Systems) or an equal volume of DMSO for 15 minutes prior to the addition of supernatants. Two hours after the addition of fresh media, 40 μL or 50 μL of culture supernatant was added to triplicate wells for 20% or 25% vol/vol concentration, respectively. The same volume of RPMI-g was added to triplicate wells as a media only control. At 5.5 or 23.5 hours after adding the supernatant, 8 μL of 10% Triton X-100 was added to three untreated wells and media was pipetted up and down to facilitate lysis of cells. Plates were returned to incubate at 37°C with 5% CO_2_ for 30 minutes, then 4 μL of media was removed from each well and added to 96 μL of LDH Storage Buffer (200 mM Trix-HCl pH 7.3, 10% glycerol, 1% BSA) in a new 96 well plate. Samples were either used immediately or stored at –80°C until use. LDH release was quantified using the LDH-Glo Cytotoxicity Assay (Promega) according to manufacturer instructions. % LDH release was calculated for each sample by standardizing against LDH values from the Triton X-100 treated wells. All cells throughout this protocol were grown or maintained at 37°C and 5% CO_2_.

### SYTOX imaging assay

Culture medium was removed from each chamber of CD14^+^ cells that were isolated and seeded in μ-Slide 8 Well chambers as described above. SYTOX Green Nucleic Acid Stain (ThermoFisher Scientific) was diluted to a concentration of 30 nM in fresh MEM-α media with M-CSF but without added FBS or penicillin/streptomycin. Media with SYTOX was added back to each chamber at a volume of 240 μL for testing supernatants at a 20% v/v concentration. 60 μL of culture supernatant was added to duplicate chambers. The same volume of RPMI-g was added to duplicate wells as a media only control. Using a BioTek Cytation 5 (Agilent), the μ-Slide was kept at 37°C and 5% CO_2_, and three randomly selected fields of view in each chamber were imaged every 30 minutes over the course of 15 hours. Both phase contrast and GFP fluorescence (excitation/emission 469/525) were imaged. Cellular analysis with cell counting was performed using Agilent BioTek Gen5 software (v. 3.16) to automatically enumerate the number of GFP+ nuclei in each image. Counts for GFP+ nuclei in the three fields of view of each well were added together, and the sum of GFP+ nuclei were averaged for replicate wells.

### SaeRS reporter assay

1 mL of *S. aureus* and *C. albicans* overnight cultures were pelleted, washed, and resuspended in 1X PBS as described for the supernatant preparation. Using RPMI-g that was generated with RPMI1640 without phenol red (Gibco), 990 μL of media was added to Eppendorf tubes for mono-cultures and 980 μL of media was added to Eppendorf tubes for co-cultures. 10 μL of *S. aureus, C. albicans,* or both organisms were added to tubes. Tubes were vortexed and 200 μL of each culture was added to triplicate wells of a black-walled, clear bottom 96 well plate. Using a BioTek Synergy HT plate reader, plate was incubated for 20 hours with orbital shaking and incubation at 37°C. Every hour, fluorescence was measured using excitation/emission of 479/520 nm to measure GFP and 579/616 nm to measure mCherry. GFP fluorescence for each SaeP reporter strain was either blanked against the isogenic *gfp* promoterless control strain or standardized against the constitutive mCherry values in each replicate culture.

### RNA isolation and quantitative RT-PCR

*S. aureus* WT and Δ*agr* were grown with or without *C. albicans* in triplicate cultures in RPMI-g medium, as described for supernatant preparation. At 6 and 12 hours of culture, organisms were pelleted and washed in 1X ice cold PBS, then pellets were frozen at –80°C until RNA isolation. RNA was isolated from each pellet as previously described (71). Briefly, frozen pellets were thawed and resuspended in 100 μL of TE Buffer. Resuspended cells were transferred to Yellow RNA Lysis Kit tubes (Next Advance) and cells were lysed for 1 min at maximum power in a Bullet Blender (Next Advance). Using the RNeasy Mini Kit (QIAGEN), 650 μL of Buffer RLT containing 1% β-mercaptoethanol was added to each tube and cells were lysed again for 1 min in the Bullet Blender. RNA was then isolated from the resulting suspension according to the manufacturer protocol. Isolated RNA was treated with the TURBO DNA Free Kit (ThermoFisher) and then 1 μg of RNA was converted to cDNA using qScript cDNA SuperMix (QuantaBio), following manufacturers’ protocols. Quantitative RT-PCR (qRT-PCR) was performed using iQ SYBR Green Supermix (Bio-Rad), cDNA, and each primer pair. Primers to quantify *saeP* transcript levels are found in (72). Primers to quantify *lukSF-PV* transcript levels are 5’-TCTCAAAACAAATGACCCCAAT-3’ and 5’-GGACCACTATTAAAATTACCACC-3’, and primers to quantify *gyrB* transcript levels are 5’-GGATCGACTTCAGAGAGAGG-3’ and 5’-CGTCCGTTATCCGTTACTTTAATC-3’. RNA from each sample without added qScript was used as a negative control for qRT-PCR. Reactions were performed using a CFX96 qPCR cycler (Bio-Rad) and following manufacturer’s guidelines with an annealing temperature of 52°C. Fold change was calculated using the ΔΔC_t_ method (73), whereby the C_t_ value for each gene was averaged from three technical replicates and standardized against *gyrB* C_t_ values from the same sample.

### DNA isolation, whole genome sequencing, and variant calling of *S. aureus* isolates

Overnight bacterial cultures of *S. aureus* were pelleted and resuspended in lysis buffer (150 mM NaCl, 25 mM Tris-HCl pH 8.0, 50 mM Glucose, 10 mM EDTA) (74). Cells were transferred to Yellow RNA Lysis Kit tubes (Next Advance) and were lysed for 1 min at maximum power in a Bullet Blender (Next Advance). Lysed samples were treated with 100 μg/mL RNase A and genomic DNA was extracted using the DNeasy Blood & Tissue Kit (QIAGEN), following manufacturer’s instructions. Isolated DNA quality and concentration were confirmed by Nanodrop and Qubit (ThermoFisher). Whole genome sequencing was performed by SeqCoast Genomics (Portsmouth, New Hampshire) using Illumina short-read sequencing to produce 150 bp paired-end reads. Raw reads were trimmed using BBDuk (75) and mapped to the reference genome *S. aureus* USA300_FPR_3757 (NCBI accession number NC_007793.1) using Geneious Prime v.2025.0.3. Variant calling was performed in Geneious for polymorphisms with a minimum 90% variant frequency and a minimum coverage of 30 reads.

### Statistical analysis

Statistical analyses were performed using Prism (GraphPad, version 10.5.0). Data were checked for normality prior to statistical analysis. To assess the effects of co-culture and strain on cytotoxicity towards host cells, a one-way ANOVA was used with a *post hoc* Tukey’s multiple comparisons test. To assess the effects of co-culture and reporter construct on fluorescence levels, a one-way ANOVA was used with a *post hoc* Tukey’s multiple comparisons test. To assess the effects of co-culture on transcript levels, a one-way ANOVA was used with a *post hoc* Dunnett’s test of multiple comparisons, to compare relative to wild-type transcript levels.

## Acknowledgements

KRE was supported by National Institute of Allergy and Infectious Diseases (NIAID) grant T32AI095202 and voucher funding from National Center for Advancing Translational Sciences CTSA award No. UL1 TR002243. JAM was supported by National Science Foundation Graduate Research Fellowship 2444112 and NIAID T32AI112541. VJT was supported by NIH NIAID grants R01AI105129 and R01AI099394, and by the American Lebanese Syrian Associated Charities (ALSAC) at St. Jude. BHP was supported by NIAID grant R01AI177615. JEC was supported by NIAID grants R01AI177615, R01AI161022, and R01AI173795. The content of this manuscript is solely the responsibility of the authors and does not necessarily represent official views of our funders or institutions.

The authors would like to thank Kimberly Davis and Irnov Irnov for providing *S. aureus* Sae reporter strains. We thank Gerry Van Horn and Jonathan Schmitz from *micro*VU and the Center for Personalized Microbiology for providing *C. albicans* clinical isolates. We also acknowledge Sandy Yoder, Buddy Creech, and the Vanderbilt Vaccine Research Program for phlebotomy and for providing *S. aureus* clinical isolates that were obtained with support from R01AI139172. Finally, we thank the members of the Cassat laboratory for providing feedback on data throughout the development of the project and for proofreading the manuscript.

## Notes

### Competing Interest Statement

The authors have declared no competing interest.

