## Supplemental Material for "*Candida albicans* activates *Staphylococcus aureus* virulence regulatory systems to drive toxin-mediated human cell death"

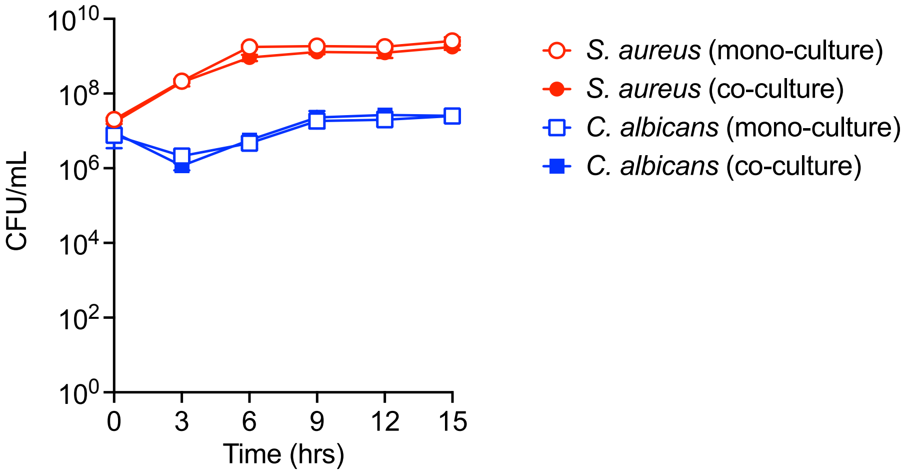


**Supplementary Figure 1. *S. aureus* and *C. albicans* growth is similar in mono-culture and co-culture.**

*S. aureus* wild-type was cultured in RPMI-g with and without *C. albicans* for 15 hours. Every 3 hours, an aliquot of culture was removed and plated on selective media to enumerate CFU/mL for *S. aureus* and *C. albicans.* Red circles represent *S. aureus* CFU/mL and blue squares represent *C. albicans* CFU/mL. Open symbols represent microbial burdens from mono-cultures and filled symbols represent microbial burdens from co-cultures. N = 3 replicates per culture and error bars represent SD. Some error bars are hidden by the symbols.

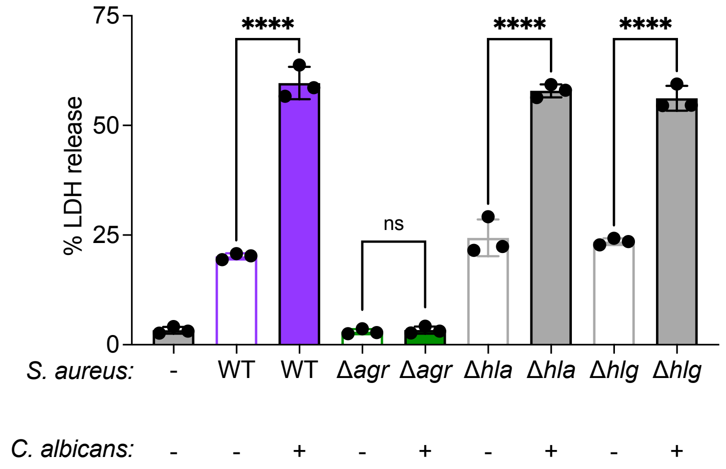


**Supplementary Figure 2. Single mutants for genes encoding α-toxin and γ-hemolysin retain cytotoxicity towards murine monocytes.**

Murine bone marrow-derived monocytes were exposed to culture supernatants from *S. aureus* strains grown with or without *C. albicans.* LDH levels in cell culture were quantified at 24 hours after supernatant exposure. % LDH release was calculated by standardizing LDH values in each well against cells lysed with 10% Triton-X 100 as a positive cell death control. *S. aureus* WT = wild-type; Δ*agr* = *agrACBD::tet*; Δ*hla* = *hla*::*spc*; Δ*hlg* = *hlgACB::tet*. ****p<0.001 and ns (not significant) by one-way ANOVA with Tukey's multiple comparisons test; n = 3 wells per supernatant condition, bars = mean±SD.

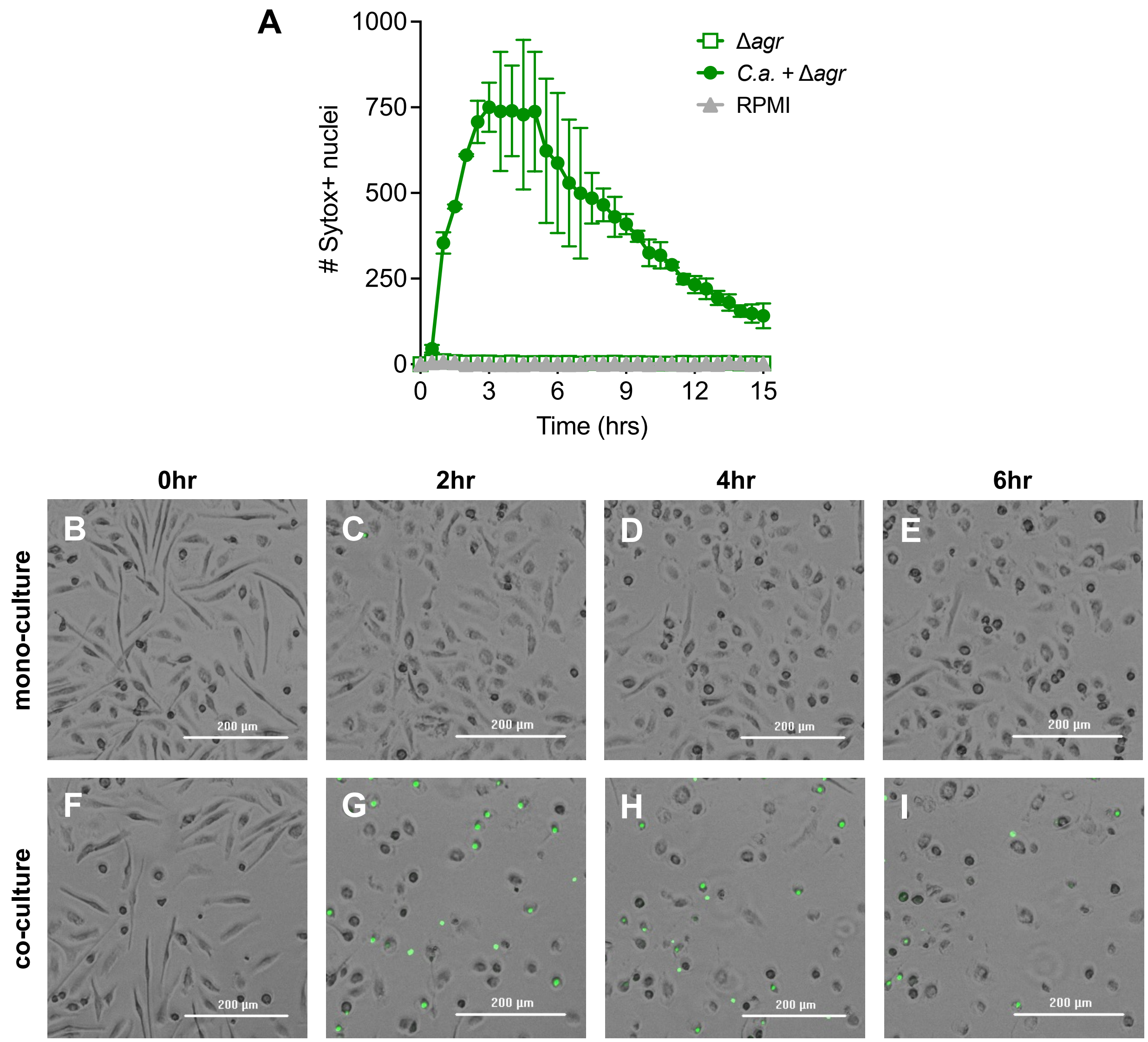


**Supplementary Figure 3. Human monocyte cell death occurs rapidly after exposure to supernatant from *S. aureus* Δ*agr* co-cultured with *C. albicans*.**

Human $CD14^+$ monocytes in media containing SYTOX Green nucleic acid stain were exposed to culture supernatants from *S. aureus* Δ*agr* grown with or without *C. albicans*. (A) SYTOX+ nuclei were enumerated in Gen5 using cellular analysis with cell counting from three randomly chosen fields of view in each well every 30 minutes over 15 hours of imaging. N = 2 replicate wells per condition, error bars represent SD. Symbols for the RPMI and Δ*agr* group are overlapping. (B-I) Representative images of monocytes at 0hr (B,F), 2hr (C,G), 4hr (D,H), and 6hr (E,I) after exposure to supernatant from *S. aureus* Δ*agr* mono-culture (B-E) or *S. aureus* Δ*agr* co-cultured with *C. albicans* (F-I).

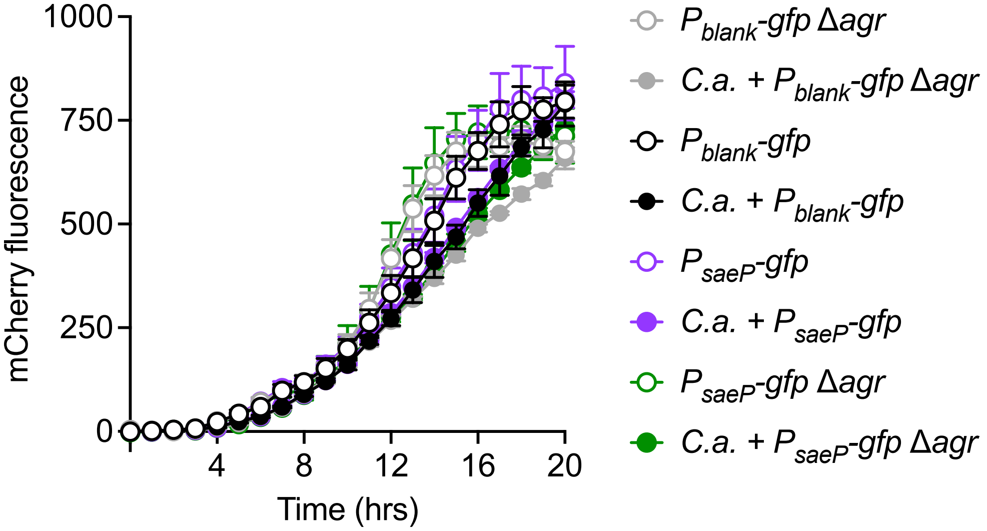


**Supplementary Figure 4. *C. albicans* co-culture does not enhance constitutive mCherry fluorescence in *S. aureus.***

*S. aureus* wild-type and Δ*agr* $P_{saeP}$-*gfp* or $P_{blank}$-*gfp* strains were cultured with and without *C. albicans* (*C.a.*), and mCherry fluorescence values were measured every hour for 20 hours. N = 3 wells or replicates per condition, error bars represent SD.

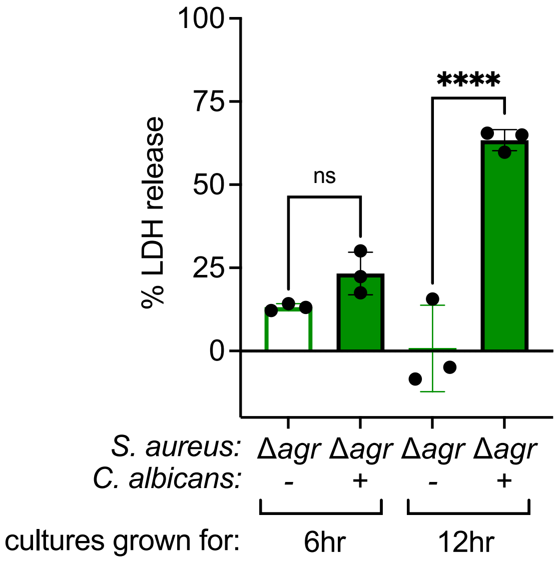


**Supplementary Figure 5. *C. albicans* induces *S. aureus* Δ*agr* cytotoxicity by 12 hours of co-culture growth.**

Human CD14$^{+}$ monocytes were exposed to supernatant from *S. aureus* strains cultured with and without *C. albicans* for either 6 hours or 12 hours in RPMI-g. % LDH release was quantified at 24 hours after supernatant exposure and calculated by standardizing LDH values in each well against cells lysed with 10% Triton-X 100 as a positive cell death control. *S. aureus* Δ*agr* = *agrACBD::tet.* ****$p<0.0001$ and ns (not significant) by one-way ANOVA with Tukey's multiple comparisons test; n = 3 wells or replicates per condition, bars = mean±SD.

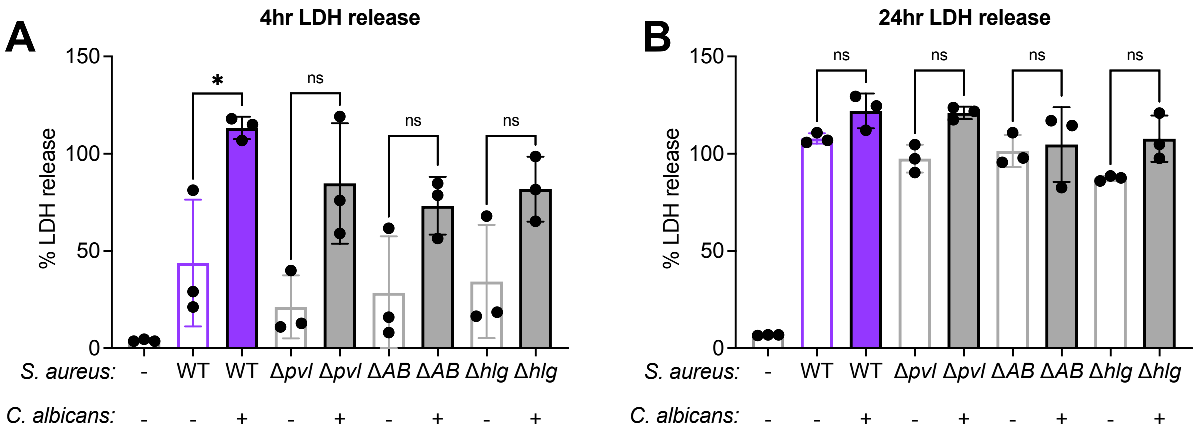


**Supplementary Figure 6. *C. albicans* co-culture enhances cytotoxicity of *S. aureus* single toxin deletion mutants towards primary human monocytes.**

Human CD14+ monocytes were exposed to supernatant from *S. aureus* strains cultured with and without *C. albicans*. % LDH release was quantified at 4 hours (A) or 24 hours (B) after supernatant exposure and calculated by standardizing LDH values in each well against cells lysed with 10% Triton-X 100 as a positive cell death control. *S. aureus* WT = wild-type; Δ*pvl* = *lukSF-PV::spc*; Δ*AB* = Δ*lukAB*; Δ*hlg* = *hlgACB::tet.* *p<0.05 and ns (not significant) by one-way ANOVA with Tukey's multiple comparisons test; n = 3 wells or replicates per condition, bars = mean±SD.

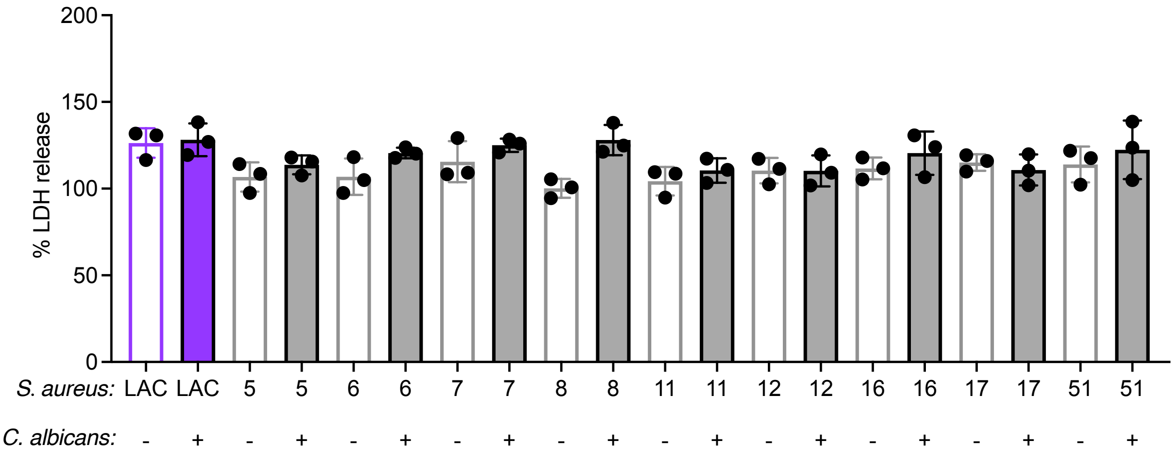


**Supplementary Figure 7. *S. aureus* LAC* and clinical isolates grown in mono-culture and co-cultured with *C. albicans* induce maximal cytotoxicity towards human cells by 24 hours.** Human CD14$^+$ monocytes were exposed to supernatant from *S. aureus* LAC* and USA300 clinical isolate strains cultured with and without *C. albicans*. % LDH release was quantified 24 hours after supernatant exposure and calculated by standardizing LDH values in each well against cells lysed with 10% Triton-X 100 as a positive cell death control.

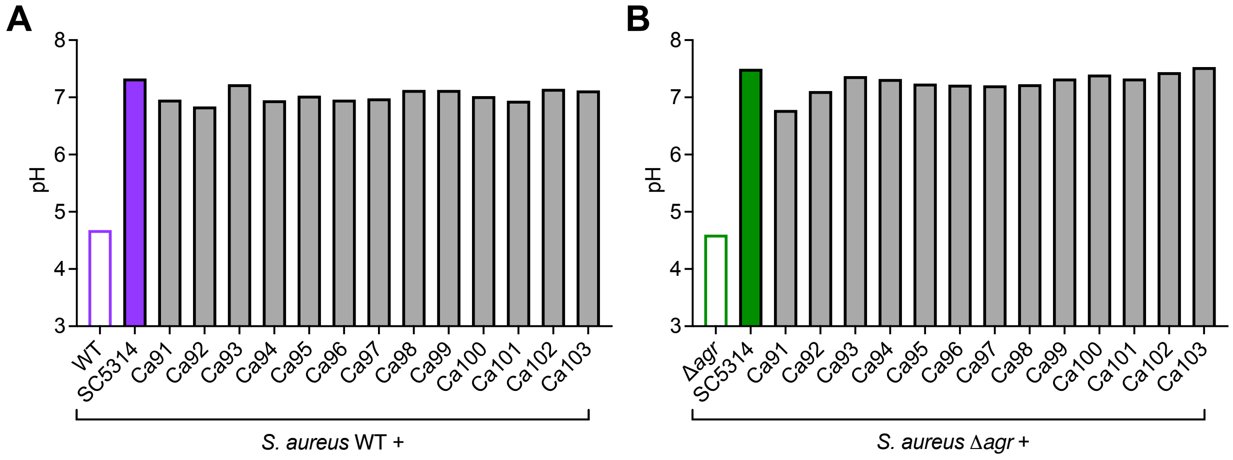


**Supplementary Figure 8. *C. albicans* SC5314 and clinical isolates maintain media at a neutral pH during co-culture with *S. aureus.***

*S. aureus* WT and Δ*agr* was cultured with and without *C. albicans* for 15 hours in RPMI-g. The media pH was measured for *S. aureus* mono-culture and in each co-culture.

**Supplementary Table 1.** *Staphylococcus aureus* strains used in this work.

| **Strain name** | **Description** | **Source** |
|---|---|---|
| AH1263 (LAC*) | Wild-type reference strain; USA300 MRSA $Erm^S$ | (1) |
| Δ*agr* | LAC* *agrACBD::tet* | (2) |
| Δ*psm* | LAC* *psmα1-4::erm* | (3) |
| Δ*hla* | LAC* *hla::spc* | (4) |
| Δ*hlg* | LAC* *hlgACB::tet* | (5) |
| Δ*psm*/Δ*hla* | LAC* *psmα1-4::erm hla::spc* | This study |
| Δ*psm*/Δ*hlg* | LAC* *psmα1-4::erm hlgACB::tet* | This study |
| Δ*psm*/Δ*hlg*/Δ*hla* | LAC* *psmα1-4::erm hlgACB::tet hla::spc* | This study |
| Δ*sae* | LAC* *sae*QRS*::spc* | (2) |
| Δ*agr*/Δ*sae* | LAC* *agrACBD::tet saeQRS::spc* | (2) |
| $P_{saeP}$-*gfp* | LAC* SapI1 *attC*::$P_{sarA}$-*sod*RBS-*mCherry* SAUSA300_RS05730::$P_{saeP}$-*gfp* | (6) |
| $P_{saeP}$-*gfp* Δ*agr* | LAC* SapI1 *attC*::$P_{sarA}$-*sod*RBS-*mCherry* SAUSA300_RS05730::$P_{saeP}$-*gfp agrACBD::tet* | This study |
| $P_{blank}$-*gfp* | LAC* SapI1 *attC*::$P_{sarA}$-*sod*RBS-*mCherry* SAUSA300_RS05730::*gfp* | (6) |
| $P_{blank}$-*gfp* Δ*agr* | LAC* SapI1 *attC*::$P_{sarA}$-*sod*RBS-*mCherry* SAUSA300_RS05730::*gfp agrACBD::tet* | This study |
| Δ*agrA* | LAC* *agrA::tn-bursa* | NARSA |
| Δ*pvl* | LAC* *lukSF-PV::spc* | (7) |
| Δ*lukAB* | LAC* Δ*lukAB* | (8) |
| Δ*agrA*/Δ*pvl* | LAC* *agrA::tn-bursa lukSF-PV::spc* | This study |
| Δ*agrA*/Δ*hlg* | LAC* *agrA::tn-bursa hlgACB::tet* | This study |
| Δ*agrA*/Δ*lukAB* | LAC* *agrA::tn-bursa* Δ*lukAB* | This study |
| Δ*agr*/Δ*pvl* | LAC* *agrACBD::tet lukSF-PV::spc* | This study |
| Δ*agr*/Δ*pvl* +*pvl* | LAC* *agrACBD::tet lukSF-PV::spc* SaPI1 *attC::*p*pvl-pvl* | This study |
| VUSA5 | USA300 ST8 clinical isolate | This study |
| VUSA6 | USA300 ST8 clinical isolate | This study |
| VUSA7 | USA300 ST8 clinical isolate | This study |
| VUSA8 | USA300 ST8 clinical isolate | This study |
| VUSA11 | USA300 ST8 clinical isolate | This study |
| VUSA12 | USA300 ST8 clinical isolate | This study |
| VUSA16 | USA300 ST8 clinical isolate | This study |
| VUSA17 | USA300 ST8 clinical isolate | This study |
| VUSA51 | USA300 ST8 clinical isolate | This study |
| VUSA5 Δ*agr* | USA300 ST8 clinical isolate *agrACBD::tet* | This study |
| VUSA6 Δ*agr* | USA300 ST8 clinical isolate *agrACBD::tet* | This study |
| VUSA7 Δ*agr* | USA300 ST8 clinical isolate *agrACBD::tet* | This study |
| VUSA8 Δ*agr* | USA300 ST8 clinical isolate *agrACBD::tet* | This study |

| | | |
|---|---|---|
| VUSA11 Δ*agr* | USA300 ST8 clinical isolate *agrACBD::tet* | This study |
| VUSA12 Δ*agr* | USA300 ST8 clinical isolate *agrACBD::tet* | This study |
| VUSA16 Δ*agr* | USA300 ST8 clinical isolate *agrACBD::tet* | This study |
| VUSA17 Δ*agr* | USA300 ST8 clinical isolate *agrACBD::tet* | This study |
| VUSA51 Δ*agr* | USA300 ST8 clinical isolate *agrACBD::tet* | This study |

**Supplementary Table 2.** *Candida albicans* strains used in this work.

| Strain name | Description | Source |
|---|---|---|
| SC5314 | Wild-type reference strain | (9) |
| Ca91 | Clinical strain isolated from the respiratory tract | This study |
| Ca92 | Clinical strain isolated from the respiratory tract | This study |
| Ca93 | Clinical strain isolated from the respiratory tract | This study |
| Ca94 | Clinical strain isolated from the respiratory tract | This study |
| Ca95 | Clinical strain isolated from the respiratory tract | This study |
| Ca96 | Clinical strain isolated from the respiratory tract | This study |
| Ca97 | Clinical strain isolated from the respiratory tract | This study |
| Ca98 | Clinical strain isolated from the respiratory tract | This study |
| Ca99 | Clinical strain isolated from the respiratory tract | This study |
| Ca100 | Clinical strain isolated from the respiratory tract | This study |
| Ca101 | Clinical strain isolated from the respiratory tract | This study |
| Ca102 | Clinical strain isolated from the respiratory tract | This study |
| Ca103 | Clinical strain isolated from the respiratory tract | This study |
